# Genome-wide CRISPRi-seq identified ferredoxin-NADP reductase FprB as a synergistic target for gallium therapy in *Pseudomonas aeruginosa*

**DOI:** 10.1101/2024.09.01.610675

**Authors:** Yu Zhang, Tingting Zhang, Xue Xiao, Adam Kawalek, Jinzhao Ou, Anmin Ren, Wenhao Sun, Vincent de Bakker, Yujie Liu, Yuelong Li, Liang Yang, Liang Ye, Ning Jia, Jan-Willem Veening, Yejun Wang, Xue Liu

## Abstract

With the rise of antibiotic-resistant bacteria, non-antibiotic therapies like gallium are increasingly gaining attention. Gallium ions exhibit potent activity against multidrug-resistant bacteria and intravenous gallium nitrite is under phase 2 clinical trials to treat chronic *Pseudomonas aeruginosa* infections in cystic fibrosis patients. However, its clinical efficacy is constrained by the achievable peak concentration in human tissue. To address this limitation, we applied a genome-wide CRISPR interference approach (CRISPRi-seq), to identify potential synergistic targets with gallium. Through the systemic screening, we classified the essential genes by response time and growth reduction, pinpointing the most vulnerable therapeutic targets in this species. In addition, we identified a highly conserved gene *fprB*, encoding a ferredoxin-NADP^+^ reductase, the downregulation of which dramatically sensitized the cells to gallium. Using a null mutant, we confirmed the loss of *fprB* lowers the minimum inhibitory concentration of gallium from 320 µM to 10 µM and shifted gallium’s mode of action from bacteriostatic to bactericidal. Further investigation revealed that FprB plays a critical role in modulating oxidative stress induced by gallium, via control of the iron homeostasis and reactive oxygen species accumulation. Deleting *fprB* also enhanced gallium’s efficacy against biofilms formation and improved outcomes in murine lung infection model of *P. aeruginosa,* suggesting FprB as a promising drug target in combination with gallium. Overall, our data showed CRISPRi-seq as a powerful tool for systematic genetic analysis of *P. aeruginosa*, advancing identification of novel therapeutic targets.

## Introduction

*Pseudomonas aeruginosa* is an opportunistic Gram-negative pathogen that frequently causes infections, particularly in immunocompromised individuals as well as in those with cystic fibrosis (CF) and chronic obstructive pulmonary disease (COPD) (Shi et al. 2023; Rossi et al. 2021; Martínez-Solano et al. 2008). It is responsible for a range of infections, including pneumonia, urinary tract infections, bloodstream infections and wound infections, which result in considerable morbidity and mortality (Qin et al. 2022). Due to its extensive resistance to carbapenems and other drugs, *P. aeruginosa* has been identified by the World Health Organization (WHO) as a critical pathogen of concern in their “ESKAPE” panel (Shi et al. 2023). Therefore, the considerable threat posed by the multi-drug resistant virulent of *P. aeruginosa* highlights the urgent need for global collaboration in developing new antimicrobial strategies.

Identifying the essential genes is a crucial stage of development of novel anti-infective agents against multidrug-resistant pathogens like *P. aeruginosa* (Juhas, Eberl, and Church 2012; Lee et al. 2015). A variety of techniques such as transposon sequencing (Tn-seq), transposon-directed insertion site sequencing (traDIS), high-throughput insertion tracking by deep sequencing (HITS) and insertion sequencing (INSeq) have been employed to globally identify essential genes through transposon mutagenesis and massively parallel sequencing in a high-throughput and comprehensive manner (Baba et al. 2006; Gallagher, Shendure, and Manoil 2011; Lee et al. 2015; Cain et al. 2020; de Bakker et al. 2022). However, these approaches have inherent technical constraints, such as requirement of large libraries to fully cover the genome due to not all transposon insertions resulting in gene inactivation, and not being able to simultaneously investigate the relative importance of essential genes under the conditions tested. In contrast, CRISPR interference (CRISPRi), which employs a catalytically inactive form of Cas9 nuclease (dCas9) in conjunction with single guide RNAs (sgRNAs) to inhibit gene expression, offers benefits over these traditional genetic methods for analyzing essential and conditionally essential gene phenotypes (Peters et al. 2016). Furthermore, quantitative CRISPRi facilitates the pinpointing of vulnerabilities in essential genes, furnishing crucial insights to fine-tune targets for drug discovery (Bosch et al. 2021). Recently, we and other groups developed CRISPRi-seq, CRISPRi in conjunction with next-generation sequencing, in multiple bacteria, and enabled genome-wide screenings to identify essential genes, antibiotic or phage sensitivity relevant factors (Bosch et al. 2021; Liu et al. 2021; Liu et al. 2023; Zhu et al. 2023; Hawkins et al. 2020; Jiang, Oikonomou, and Tavazoie 2020; Wang et al. 2018; Rousset et al. 2018; Donati et al. 2021). However, a genome-wide CRISPRi-seq toolbox to study *P. aeruginosa* gene fitness is still lacking. This gap is likely attributable to the challenges in developing a tightly regulated inducible system, which is crucial for the efficient and uniform expression of sgRNAs targeting essential genes. The commonly employed arabinose-inducible system in *P. aeruginosa*, for instance, suffers from leakage (Qu et al. 2019), resulting in basal CRISPRi activity. Additionally, the lack of an effective strategy for cloning sgRNAs has further hindered the application of genome-wide CRISPRi in *P. aeruginosa*. In this study, we addressed these challenges by establishing a rigorously controlled tetracycline (tet)-inducible CRISPRi system in *P. aeruginosa*, demonstrating its efficiency and tunability across a wide range of strains, including both standard laboratory and clinically relevant isolates. Furthermore, we employed a *ccdB*-based counter selection system for efficient sgRNA cloning, and successfully developed a comprehensive genome-wide CRISPRi library targeting 98% of the genetic elements in the *P. aeruginosa* PA14 strain.

Gallium has been approved by the U.S. Food and Drug Administration (FDA) as a therapy for cancer-related hypercalcemia (Leyland-Jones 2004), and was also recognized as a promising antibacterial agent (Chatterjee et al. 2016; Kaneko et al. 2007). Recently, a proof-of-principle phase 1 clinical trial of gallium in CF patients with chronic *P. aeruginosa* lung infections proved its safety and efficacy (Goss et al. 2018). This phase 1 clinical study used intravenous gallium nitrate which was already approved by the U.S. FDA for hypercalcemia of malignancy. A follow-up Phase II clinical trial in 2016 (NCT02354859) showed a marked reduction in *P. aeruginosa* in the sputum of CF patients receiving gallium nitrate versus the placebo cohort, despite not meeting the primary endpoint (Reig, Le Gouellec, and Bleves 2022). The efficacy of gallium therapy for *P. aeruginosa* CF patients is limited by the peak plasma and sputum concentrations, which is 8 ~ 12 µM (Goss et al. 2018). Gallium, which has an ionic radius almost identical to that of iron, can be taken up by bacteria in place of iron (Chitambar and Narasimhan 1991). Once inside the cell, gallium is incorporated into iron-containing proteins but cannot substitute for iron in its essential biological functions, such as redox cycling. In *P. aeruginosa*, loss-of-function mutations that confer increased tolerance to gallium therapy have been identified. Notably, HitA and HitB, which comprise the major transporters of iron, have emerged as key factors in this process (Goss et al. 2018). However, the genes potentially associated with increased sensitivity upon loss of function remain largely unexplored, impeding the advancement of synergistic strategies to enhance the efficacy of gallium treatment. To address this, we successfully used genome-wide CRISPRi library and screened gallium related factors.

Using the CRISPRi library, we developed and implemented quantitative CRISPRi-seq to refine the essential gene list and assess the vulnerability of essential genes in *P. aeruginosa* PA14. Furthermore, we conducted CRISPRi-seq to pinpoint genes associated with sensitivity to gallium. The analysis revealed 98 genes whose repression results in increased tolerance to gallium and 9 genes whose repression leads to heightened sensitivity. This provides the first comprehensive genome-wide overview of genes involved in gallium sensitivity. The identification of *hitAB* as the foremost genes associated with increased gallium tolerance following CRISPRi knockdown reaffirms findings from prior studies (Goss et al. 2018; Guo et al. 2019; Zemke et al. 2020; García-Contreras et al. 2013), underscoring the effectiveness of CRISPRi-seq in elucidating mechanisms within *P. aeruginosa*. From the screening results, we selected the 11 top candidates for further validation, which confirmed 8 out of the 11 genes, further demonstrating the robustness and utility of CRISPRi-based chemical genetic studies in *P. aeruginosa*. Remarkably, CRISPRi-seq unveiled the significance of a ferredoxin-NADP^+^ reductase (Fpr)-coding gene, *fprB*, in conferring resistance to gallium. Loss-of-function mutations in FprB resulted in approximately a 30-fold increase in *P. aeruginosa*’s susceptibility to gallium in terms of MIC. Phylogenetic analysis revealed that FprB is a remarkably conserved protein in *P. aeruginosa*, present in over 99% of the strains with whole genome sequences, whereas lacking homology in human. The effect of loss-function of FprB for gallium sensitivity was reaffirmed by targeting biofilm formation and with murine lung infection models. Clinical studies have demonstrated that the peak plasma and sputum concentrations of gallium achieved through FDA-approved intravenous gallium nitrate treatment typically range from 2 to 3 μg/mL, equivalent to approximately 8 to 12 μM (Goss et al. 2018). Notably, these concentrations remain relatively stable and cannot be enhanced through increased dosage (Goss et al. 2018), underscoring the importance of identifying synergistic targets for gallium treatment, such as the identified FprB. Efforts are underway to identify effective inhibitors for FprB and evaluate their efficacy in enhancing gallium therapy for CF patients infected with *P. aeruginosa*, especially the multidrug resistant strains.

## Materials and Methods

### Bacterial strains and growth conditions

All strains and plasmids utilized in this study are enumerated in **Supplementary Table S1**, while primers are in **Supplementary Table S2**. *Pseudomonas* strains used in this study includes *P. aeruginosa* PAO1, PA14, P5 and PDO300. Routine cultures for *Pseudomonas* strains were performed at 37_ in Luria-Bertani (LB) broth or on LB agar plates with antibiotic where necessary: 20 μg/mL gentamicin and 30 μg/mL apramycin. For gallium treatment CRISPRi-seq screen, PA14 were cultured in M9 minimal medium containing Na_2_HPO_4_·7H_2_O (47.8 mM), KH_2_PO_4_ (22 mM), NaCl (8.6 mM), NH_4_Cl (18.6 mM), MgSO_4_ (1.0 mM), CaCl_2_ (0.1 mM), and sodium citrate (20_mM) as the carbon source at 37°C under vigorous aeration. *E. coli* WM3064 represents a diaminopimelic acid (DAP)-auxotrophic strain (Wang et al. 2015), offering significant utility for conjugation experiments and replication of plasmids necessitating Pir protein. *E. coli* strains were cultivated in LB broth at 37°C, supplemented with antibiotics where necessary: 20 μg/mL gentamicin and 100 μg/mL ampicillin. Cultures were subjected to CRISPRi repression, primarily induced with 25 ng/mL doxycycline (Dox), unless stated differently.

### Development of a tetracycline (tet)-inducible platform in *P. aeruginosa*

The tet-inducible system for *P. aeruginosa* was developed based on our previously published tet-inducible system in *Streptococcus pneumoniae*, which was proved to be very tightly controlled (Sorg et al. 2020; Liu et al. 2021). A gblock (gblock 6-TetPt5) containing *Pseudomonas* codon-optimized *tetR* (termed *tetR-opt*), a constitutive P*_con_* promoter and tet-inducible P*_tet_* promoter were synthesized by Sangon (Sangon, China). The sequence of gblock 6-TetPt5 is shown in **Supplementary Table S3.** The tet-inducible system uses a modified pUC18T-mini-Tn7T-P3-luxABCDE-based integration vector (**Supplementary Table S1**), incorporating a constitutively expressed *tetR-opt* driven by the P*_con_* promoter, amplified with primers OXL384 and OXL567 (**Supplementary Table S2**). The P*_3_* promoter was substituted with a P*_tet_* (amplified with OXL387 and OXL568) to drive *luxABCDE* reporter gene, enabling precise quantification of tet-regulated gene expression. Herein, we designated the plasmid containing the tet-inducible system as Ptet-lux-Tn7T-Gm. All PCR fragments were amplified using Phanta DNA Polymerase (Vazyme, P515-03, China) in preparation for In-Fusion cloning with the ClonExpress Ultra One Step Cloning Kit V2 (Vazyme, C116-02, China).

### Construction of the *ccdB* counterselection system for sgRNA cloning

The *ccdB* counterselection system (Bernard 1996) was employed to enhance the efficiency of sgRNA cloning through negative selection against non-modified plasmids. Firstly, *E. coli* WM3064 with mutated *gyrA* (462_Arg→Cys_) was engineered via CRISPR/Cas9 editing as previously reported (Chen et al. 2018), to confer resistance to the CcdB toxin (Bernard and Couturier 1992). Sequences of the sgRNA and single strand DNA repair template designed to introduce the gyrA462_Arg→Cys_ mutation by CRISPR/Cas9 editing were listed in the oligo list (**Supplementary Table S2**). The obtained *E. coli* WM3064 gyrA462_Arg→Cys_ was confirmed by Sanger sequencing, used as the host for construction of sgRNA cloning vector with *ccdB* counter selection system. The *ccdB* gene flanked by two BsaI sites, ordered as the gBlock “gblock4-ccdB-CSY” (**Supplementary Table S3**) was amplified using primers OXL289 and OXL290, and then cloned into the Mobile-CRISPRi plasmid pJMP2846 (Addgene#160676) (Banta et al. 2020) via BsaI-mediated Golden Gate Assembly. The product of Golden Gate Assembly was transformed into *E. coli* WM3064 gyrA462_Arg→Cys_ followed by selection with 30 μg/mL apramycin on LB agar plates supplied with 300 μM DAP. The resulting plasmid is named pJMP2846-*ccdB*, which serves as the vector for constructing the CRISPRi system and subsequent sgRNA cloning.

### Construction of the tet-inducible CRISPRi system

Our CRISPRi system was primarily constructed based on pJMP2846-*ccdB*. The tet-inducible element mentioned above was amplified and introduced to replace the native IPTG-inducible part (*PLlacO1-LacI)* that drives dCas9 expression in pJMP2846-*ccdB*, using Primers OXL657, OXL658, OXL659 and OXL660 (**Supplementary Table S2**). The cassette containing Illumina read 1, P_3_ promoter, BsaI site-flanked *mCherry*, sgRNA scaffold, sgRNA terminator, and Illumina read 2 was generated from the plasmid pPEPZ-sgRNAclone (Addgene#141090) (de Bakker et al. 2022) using primers OXL226 and OXL227. The kanamycin resistance gene segment was replaced with one conferring apramycin resistance gene using primers OXL483 and OXL484. The construct was then transformed into *E. coli* WM3064 and selected with 30 μg/mL apramycin on LB agar plate supplied with 300 μM DAP. The resulting plasmid was named as pCRISPRi-*ccdB*. To test the efficiency of pCRISPRi-*ccdB* for sgRNA cloning and counter selection, sgRNA targeting different genes were cloned by BsaI mediated Golden Gate Assembly. Two 24-nt oligonucleotides with 5’-ends of TATA and AAAC, respectively, were synthesized for each guide sequence and annealed in TEN buffer (10 mM Tris, 1 mM EDTA, 100 mM NaCl, pH 8.0) in a thermocycler. The annealing process involved incubation at 95°C for 5 minutes and then slowly cooling them to room temperature to allow for proper pairing. The annealed oligonucleotides and pCRISPRi with *ccdB* vector were assembled based on the One-step Golden Gate protocol (Chen et al. 2018), using DNA restriction enzyme BsaI (Lablead, F5518S, China) and ligase T4 (Vazyme, C301-01, China). The assembly mixtures were transformed into *E. coli* WM3064 and plated on LB agar plates containing 30 μg/mL apramycin and 300 μM DAP for selection of successful assembly clones. As a control to test the efficiency of CcdB counter selection, the intact pCRISPRi-*ccdB* plasmid without sgRNA cloning was transformed into *E. coli* WM3064 in parallel.

### Construction of pooled CRISPRi library in *P. aeruginosa* PA14

#### sgRNA design

The sgRNAs were designed by our previously published R script for automatic sgRNA selection (https://github.com/veeninglab/CRISPRi-seq) (de Bakker et al. 2022). All sgRNAs were designed to target the coding region of the gene. For each gene, 2 sgRNAs were designed and assigned into two sets. The best sgRNA hits were assigned as set 1, and the 2^nd^ best hit as set 2. For the genes with less than 2 PAMs in the coding regions, only one sgRNA for set 1 was designed. To make the genome-wide sgRNA pool, 5981 sgRNAs were designed in set 1 sgRNA pool and 5971 sgRNAs were designed in set 2 sgRNA pool. The 20-nt spacer sequence of the designed sgRNAs and their targets were listed in **Supplementary Table S4**.

#### sgRNA cloning

The set 1 and set 2 pooled sgRNA oligos were synthesized as one oligo chip by GenScript (Nanjing, China), and the oligo sequences were listed in **Supplementary Table S4**. Each oligo contains a 20-nt sgRNA spacer sequence flanked with 2 BsaI sites, the set specific sequence for amplification of either set 1 or set 2 sgRNAs, and the universal sequence for amplification of all the sgRNAs in one chip. Set 1 and set 2 sgRNA pools were amplified with primer pairs OXL409/OXL410 and OXL411 /OXL412 (**Supplementary Table S2**) by PCR, respectively. The amplicons were purified with a Monarch DNA Cleanup kit (NEB, T1030S, USA). The purified amplicons were cloned into the plasmid pCRISPRi-*ccdB* by BsaI mediated Golden Gate Assembly. Specifically, for each Golden Gate Assembly reaction, 300 ng of the amplicons were mixed with 500 ng of the pCRISPRi-*ccdB* plasmid in a 200-μL PCR tube, followed by addition of 1 μL of with BsaI (Lablead, F5518S, China), 1 μL of T4 ligase (Vazyme, C301-01, China) and 1 µL of 10× T4 ligation buffer. MiliQ water was added into the PCR tube to bring the final volume to 20 µL. The reaction mixture was incubated in a thermocycler with the following program: 150 rounds of 37_ for 1.5 min, 16_ for 3 min, and 1 round of 37_ for 5 min, 80_ for 10 min. The product of the Golden Gate Assembly reaction was then transformed into chemical competent *E. coli* WM3064 and the success transformants were selected on LB agar plates with 30 μg/mL apramycin and 300 μM DAP on LB agar plates. In total 60 of 20-uL Golden Gate Assembly reaction products were performed and transformed. Over 250,000 transformant colonies were harvested for each sgRNA pool, indicating more than 50 times sgRNA library coverage.

#### Construction the CRISPRi library in P. aeruginosa PA14 via triparental conjugation

The *E. coli* WM3064 with sgRNA library were served as donor strain and subsequently conjugated into *P. aeruginosa* PA14 strain with a helper WM3064 strain harboring a plasmid (pJMP1039, Addgene#119239) that expresses TnsABCD transposase. The triparental conjugation was performed as describe previously (Banta et al. 2020). The successful transconjugants with integration of Tn7 cassette into PA14 chromosome were selected on LB agar plates supplemented with 30 μg/mL apramycin at 37°C. In total, the CRISPRi library construction involved collecting around 300,000 colonies for the set 1 sgRNA pool and 570,000 colonies for the set 2 sgRNA pool.

### CRISPRi-seq Screen

#### Sample preparations and sequencing

The gene essentiality screen of *P. aeruginosa* PA14 was conducted over approximately 21 generations of growth in triplicate, and samples of 7 generations, 14 generations and 21 generations were collected and analyzed by Illumina sequencing. Pooled libraries were cultured in LB broth at 37°C until reaching an optical density of 0.6 at 600 nm. The cultures were then diluted 1:100 into fresh LB broth with or without the inducer Dox (25 ng/mL). Upon reaching an OD600 of 0.6 again, 5 mL of bacteria culture was collected as the samples of 7 generations. Consecutively, the cultures underwent two more 1:100 dilutions into fresh LB broth to allow for 14 and 21 generations of Dox induction followed by bacterial pellet collection. Bacterial pellets harvested at 7, 14 and 21 generations were utilized for genomic DNA isolation using a FastPure Bacteria DNA Isolation Mini Kit (Vazyme, DC103-01, China) according to the manufacturer’s protocol. For CRISPRi-seq screen under Ga(NO_3_)_3_ treatment, pooled libraries were pre-induced in LB broth with Dox for 14 generations, followed by treating with 200 μM Ga(NO_3_)_3_ plus Dox in M9 medium and the culture continued until 21 generations. Guide encompassing region was amplified from genomic DNAs using a one-step PCR process, which incorporated Illumina barcodes N701-N712 and N501-N512 as index 1 and index 2, respectively, yielding 303 bp products (de Bakker et al. 2022). The amplicons were sequenced on an Illumina NovaSeq system by Haplox (Shenzhen, China) following the manufacturer’s protocol.

### CRISPRi-seq data analysis

The absolute abundance of each sgRNA per condition from the raw paired-end sequencing data of the CRISPRi-seq screen is determined with 2FAST2Q (v2.5.0) (Bravo, Typas, and Veening 2022). Using the default configuration of 2FAST2Q, a nucleotide-based quality filtering step discards all trimmed reads with a Phred score below 30. The depletion or enrichment of sgRNAs was analyzed using DESeq2 in R (version 4.3.1) (Love, Huber, and Anders 2014). In the gene essentiality screen, fitness values were calculated by comparing the CRISPRi induction with Dox to the non-induction condition (+Dox Vs −Dox). Gene essentiality was determined by evaluating the log_2_(Fold change, FC) values from both set 1 and set 2 screening results over 7, 14, and 21 generations. Essential genes were identified for those with log_2_FC_<_−1 and adjusted *P*-value < 0.05. The results from individual analysis of set 1 and set 2 across 7, 14 and 21 generations were displayed in **Supplementary Table S5**. In the gene categorization, genes were firstly classified into 3 categories, including “nonessential” “vulnerable” and “invulnerable” according to the standard demonstrated in **Extended Fig. 2c**. The set with the bigger |log_2_FC| was chosen to classify each gene’s essentiality. Within the vulnerable category, genes were further divided into quick- and slow-responsive, based on whether the 7-generation log_2_FC was less than −2 or greater than −2, respectively. Similarly, within the invulnerable group, genes were classified as quick responders if the 7-generation log_2_FC was less than −1, and slow responders if it was more than or equal to −1. For genes identified as nonessential, the screening results from set 1 were used as the definitive reference. The essential gene list was compared with three Tn-seq studies conducted in *P. aeruginosa* PA14 (Liberati et al. 2006; Skurnik et al. 2013; Poulsen et al. 2019). Core essential genes identified by Poulsen et al. (Poulsen et al. 2019) were highlighted in the list. Operon information was extracted from the study by Wurtzel et al. (Wurtzel et al. 2012). The generated results, including the classification of genes into five categories, comparison with Tn-seq studies, and operon information, are presented in **Supplementary Table S6**.

For the gallium treatment CRISPRi-seq screen, fitness values were determined by comparing the Ga(NO_3_)_3_-treated condition with Dox to the untreated condition [+Ga(NO_3_)_3_ Vs −Ga(NO_3_)_3_]. sgRNA targets were classified as up-regulated genes if their fitness showed a log_2_FC > 2 with an adjusted *P*-value < 0.05, and as down-regulated if the log_2_FC was less than −2 with the same *P*-value threshold. The results are provided in **Supplementary Table S7**. GO enrichment analysis for the up- and down-regulated targets was conducted and is presented in **Supplementary Table S8**.

### Construction of knockdown and knockout strains

The gene knockdown strains were engineered using a method akin to library construction but utilized a singular, targeted sequence instead of a diverse sequence pool. The specific gene knockdowns, along with the corresponding primers and bacterial strains, are presented in the results section or supplementary materials. The genes of interest in *P. aeruginosa* PAO1 were deleted via CRISPR-Cas9 genome editing with pCasPA/pACRISPR system as previously described (Chen et al. 2018). In brief, the pCasPA plasmid expressing Cas9 was first introduced and selected with 100 μg/mL tetracycline. The pACRISPR plasmid containing sgRNA was then electroporated with homologous repair templates flanking the target gene into PAO1 cells harboring pCasPA. The homologous regions were PCR amplified and then purified through gel extraction. Successful knockout mutants were isolated under 100 μg/mL tetracycline and 200 μg/mL carbenicillin selection and validated by sequencing. To achieve gene complementation, the ptet-lux-Tn7T-Gm plasmid, which facilitates gene expression under a tet-inducible promoter, was utilized. The gene’s open reading frame was PCR-amplified from PAO1 genomic DNA using primers with 15 bp extensions and subsequently cloned into the ptet-lux-Tn7T-Gm vector through In-Fusion cloning technology (Vazyme, China). This recombinant plasmid was introduced into the WM3064 strain via transformation and selected using 20 mg/mL gentamicin and 300 μM DAP. Following triparental conjugation, the fragment with gene complementation was incorporated into the genome of the PAO1 knockout strain. Complementary mutants were selected on LB agar containing 20 μg/mL gentamicin and further confirmed by PCR and sequencing.

### Growth and luminescence measurements

For each strain examined, working stocks were revived and diluted 1:100 in fresh LB broth with or without varying concentrations of Ga(NO_3_)_3_ or Dox, to initiate the cultures. The OD600 was standardized to 0.003 for all strains. A total of 100 µL of the bacterial culture were added into 96-well flat-bottom cell culture plates with 3 replicates (NEST Biotechnology, 701011, China) and employed a Tecan Spark microplate reader (Tecan, Switzerland) at 37°C for periodic monitoring of bacterial growth, with an interval time of 10 minutes. For the detection of *luxABCDE* bioluminescent signals, we transferred the cultures to 96-well flat clear bottom black microplates (αPLUS, WP96-48CMED, China) and recorded measurements at 10-minute intervals at 37°C using the Tecan Spark.

### Swarming motility assays

Swarming motility assays were conducted based on established methods with specific modifications (Kollaran et al. 2019). The swarming medium was composed of 0.5% LB agar supplemented with 100 µM CaCl_2_, 2 mM MgSO_4_, and 0.4% glucose. Each plate was centrally inoculated with 2 µL of bacterial suspension in LB broth, with the culture’s OD600 adjusted to 0.2. Plates were then incubated at 37°C for overnight.

### Determination of Minimum Inhibition Concentration of Ga(NO_3_)_3_

Minimum inhibitory concentrations (MICs) of Ga(NO_3_)_3_ were determined using the broth microdilution assay in 96-well polystyrene plates (NEST Biotechnology, 701011, China) as previously described (Dousa et al. 2024). Bacterial suspensions, adjusted to an OD600 of 0.0015, were inoculated into Mueller-Hinton broth (Oxioid, CM0405, USA) containing Ga(NO_3_)_3_ at concentrations ranging from 0 to 2560 μM (Oxoid, CM0405, USA). The plates were incubated at 37 °C for 24 hours, followed by visual inspection and OD600 measurement using Tecan Spark.

### Gallium time-killing assay

For gallium killing assays, bacterial cultures (OD600 = 0.5 ~ 0.7) were diluted to an OD600 of 0.003 in LB broth and incubated at 37°C, with or without Ga(NO_3_)_3_ supplementation. An aliquot was plated to determine the colony-forming units (CFU) at Time 0, prior to the addition of Ga(NO_3_)_3_. At specified time points, aliquots were taken and washed with sterile PBS. The cells were then serially diluted and plated on LB agar to quantify the surviving bacteria.

### Intracellular gallium and iron profiling

The methodology for assessing gallium levels in PAO1 strains, including PAO1Δ*fprB* and complemented strain, was based on the previously established protocol (Tovar-Garcia et al. 2020). Bacterial cultures (OD600 = 0.003) were grown in 6 mL of LB broth, either with or without 10 µM Ga(NO_3_)_3_, for 16 hours, and then washed twice using 50 mM HEPES buffer (pH 7.2). The samples were resuspended in 50 mM HEPES buffer and adjusted to an OD600 of 0.6, followed by centrifugation at 8000 *g* for 5 minutes to pellet the cells. Lysis was performed using 70% nitric acid at 100°C for 1 hour. Sample volumes were then normalized to 2 mL and determined by inductively coupled plasma mass spectrometry (ICP-MS) with a NexION 300X instrument (PerkinElmer, USA). To detect intracellular iron, bacterial cultures (20 mL, initial OD600 = 0.05) were cultivated with or without Ga(NO_3_)_3_ at concentrations of 12.5 and 50 µM, and harvested after 16 hours of incubation. These cultures were then sonicated at 200W for 1 minute. Subsequently, cell lysates were prepared for intracellular iron content analysis, employing a previously established method with slight modifications (Zhong et al. 2023). The Total Iron Colorimetric Assay Kit (E-BC-K772-M, Elabscience, China) was used to measure the total iron content in bacteria, and the Ferrous Iron Colorimetric Assay Kit (E-BC-K773-M, Elabscience, China) was utilized to determine the levels of intracellular ferrous iron (Fe^2+^) using a microplate reader (Tecan, Switzerland) at 593 nm.

### Quantification of intracellular ROS and H_2_O_2_ by flow cytometry

Carboxy-H2DCFDA and PeroxyOrange-1 (Thermo Fisher Scientific, USA) fluorescent probes were employed for detection of total intracellular reactive oxygen species (ROS) and hydrogen peroxide (H_2_O_2_), respectively. Upon oxidation, these probes generate fluorescent signals that can be quantified by flow cytometry, thereby indicating the extent of oxidative stress within cells (Hong et al. 2017). Bacterial cultures (OD600 = 0.5 ~ 0.7) were diluted to an OD600 of 0.01 in LB broth, either in the absence or presence of Ga(NO_3_)_3_, and incubated for 4 hours at 37°C. Following this incubation, Carboxy-H2DCFDA (final concentration of 10 μM) or PeroxyOrange-1 (final concentration of 5 μM) was added to the growth medium, and the cultures were further incubated in the dark for 2 hours at 37°C with shaking at 250 rpm. An unstained control sample was included to assess the background fluorescence. For each sample, a total of 100,000 ungated events were recorded for each sample. The NovoCyte Advanteon Flow Cytometer (Agilent, USA) was used to measure carboxy-H2DCFDA fluorescence, while the FACSAria II Flow Cytometer (BD Biosciences, USA) was employed for Peroxy Orange-1 fluorescence detection. Data analysis was conducted utilizing FlowJo software version 10 (BD Biosciences).

### NADP^+^ and NADPH detection

Total oxidized and reduced nicotinamide adenine dinucleotide phosphates (NADP^+^ and NADPH) were quantified using the NADP/NADPH-Glo™ Assay (Promega, G9081, UK) according to the manufacturer’s protocol. Bacterial cultures were diluted to an OD600 of 0.01 and treated with or without Ga(NO_3_)_3_ for 4 hours at 37°C with shaking at 250 rpm. The bacteria were harvested by centrifugation at 7,000 *g* for 2 minutes and washed with distilled water at 4°C. To ensure complete cell lysis, an equal volume of 0.2 N NaOH containing 1% DTAB (Sangon, A600431, China) was added. Following the manufacturer’s guidelines, NADP/NADPH-Glo™ Detection Reagent was introduced, and luminescence was measured using a Tecan Spark microplate reader.

### Protein purification of FprB

The *fprB* coding sequence was cloned into the pGood_6p vector and transformed into *E. coli* Rosetta (DE3) cells, allowing expression of a GST-tagged protein. Single colony was picked and cultured overnight in LB with 100 µg/mL ampicillin at 37°C and 220 rpm, then diluted 1:100 into 200 mL of LB broth and grown to an OD600 of 0.8. Protein expression was induced with 1 mM IPTG, and cultures were incubated overnight at 16°C with shaking 220 rpm. Cells were harvested by centrifugation, resuspended in buffer (20 mM Tris-HCl, 150 mM NaCl, pH 8.0), and lysed by sonication. The lysate was centrifuged, and the supernatant was incubated with Glutathione Sepharose 4B (Cytiva, 17075605, USA) beads for 4 hours at 4°C. After washing with buffer (20 mM Tris-HCl, 1 M NaCl, pH 8.0), protein was eluted with buffer (20 mM Tris-HCl, 150 mM NaCl, 20 mM GSSH). The GST tag was removed using P3c protease. FprB protein was further purified by size-exclusion chromatography and stored in buffer (20 mM Tris-HCl, 150 mM NaCl, pH 8.0, 20% glycerol).

### FprB enzyme activity assay in vitro

FprB enzyme activity was assessed using an NADPH-dependent 2,6-dichlorophenolindophenol (DCPIP, from Biosynth, FD32424, Switzerland) reduction assay as previously described (Martinez-Julvez et al. 2017). Briefly, a reaction mixture was prepared in a clear 96-well microplate, containing 800 nM FprB, 400 μM NADPH, 800 µM DCPIP, 2% DMSO, and 50 mM Tris/HCl buffer (pH 8.0). Absorbance at 620 nm was measured using a Tecan Infinite M100PRO reader at 25°C. Negative control wells contained the reaction mixture without FprB. Each condition was tested in triplicate.

### RNA-Sequencing

*P. aeruginosa* strains (PAO1 and Δ*fprB*) at an OD600 of 0.5 ~ 0.7 were harvested and rinsed twice with nuclease-free water. Total RNA was isolated using a RNAprep Pure Cell/Bacteria Kit (TIANGEN, DP430, China). Transcriptome sequencing was performed on an Illumina NovaSeq 6000 platform (Illumina, USA) by Novogene Technology Co., Ltd. (Beijing, China). Differential expression between the two groups was analyzed using the DESeq2 package. Genes with a Benjamini-Hochberg-adjusted *P*-value < 0.05 and |log_2_FC| > 2 were classified as differentially expressed.

### Distribution, evolution analysis and sequence comparison of FprB

#### Distribution analysis

Full chromosome sequences of 981 *P. aeruginosa* strains were downloaded from the NCBI Genome database (https://www.ncbi.nlm.nih.gov/genome, assessed [03/04/2024]). To detect FprB orthologues, the *P. aeruginosa* PAO1 FprB (PA4615) and its protein sequence (NP_253305.1) were used as the queries. To detect the FprB in the *P. aeruginosa* genomes, the sequence of NP_253305.1 was used to align against with the peptide sequences derived from the genome sequences of the *P. aeruginosa* strains. BLASTP was used for the sequence alignment. The criterion for homology was set as ‘coverage * identity’ >= 0.6.

#### Gene collinearity analysis

The nucleotide sequence containing the *fprB* gene and its flanking sequences, that is, 5-kb for the upstream and downstream of the center of *fprB* gene respectively, was retrieved from the genome of each *P. aeruginosa* strain. PipMaker was used to make the collinearity analysis between each pair of the sequences (http://pipmaker.bx.psu.edu/pipmaker/) (Schwartz et al. 2000).

#### Multiple sequence alignment and phylogenetic analysis

CLUSTALW was used for FprB protein sequence alignment (https://www.ebi.ac.uk/jdispatcher/msa/clustalo) (Madeira et al. 2024). The FprB multiple sequence alignment results were used for phylogenetic analysis using MEGA 7.0 with the Neigbor-Joining algorithm (https://www.megasoftware.net/) (Kumar, Stecher, and Tamura 2016).

### Biofilm assays

Biofilm quantification was evaluated in 96-well microtiter plates following established protocols (Haney, Trimble, and Hancock 2021). *P. aeruginosa* cultures were grown to an OD600 of 0.6. The cultures were then diluted 1:200 in LB broth. Subsequently, 100 μL of the diluted cultures were added to each well, with triplicates for each condition. Following a 24-hour incubation, the media supernatant was discarded, and the wells were washed with distilled water. Biofilms were then fixed with 95% methanol for 15 minutes and stained with 0.5% crystal violet for 10 minutes. The wells were rinsed again with distilled water, and 75% ethanol was added to dissolve the dye. Absorbance was measured at 590 nm after incubation.

### Mouse infections

The study protocol of animal experiments was approved by the Institutional Animal Care and Use Committee of Shenzhen University Medical School (Approval Number: IACUC-202400111). Female BALB/c mice (18 ~ 22 g, 6 ~ 8 weeks old) were purchased from Guangdong Medical Laboratory Animal Center and maintained under specific pathogen-free conditions. Prior to infection, mice were randomly assigned to either treatment or control groups. Mice were anesthetized and intranasally inoculated with 10^7^ CFU in 40 µL of cell suspension. After 3 hours, mice were treated intraperitoneally with either 50 µL PBS or 250 mM Ga(NO_3_)_3_. At 27 hours post-infection (h.p.i.), mice were necropsied and lungs were collected for bacteriological analysis. Lung tissue samples were homogenized, serially diluted in sterile PBS, and plated onto LB agar to enumerate bacterial load.

### Statistical analysis

GraphPad Prism (version 8.4.3) and R (version 4.3.1) were used for all statistical analyses. The experiments were conducted in at least triplicate to ensure biological reproducibility. Reported values represent the mean and standard deviation (SD) across the three replicates. Statistical significance between groups was determined by Two-way-measure of variance (ANOVA) and the Turkey’s multiple comparisons test at a p-value threshold of 0.05. The illustrative diagrams were created or generated using Adobe Illustrator 2021 or BioRender (https://www.biorender.com/).

### Resource availability

#### Materials availability

The vectors pCRISPRi-*ccdB* and ptet-lux-Tn7T-Gm were deposited at Addgene (catalog #226432 and 226434). The pooled CRISPRi sgRNA libraries (set 1 and set 2) for *P. aeruginosa* PA14 will be available through Addgene (#xxxxx, under processing).

#### Data availability

All sequenced data supporting the findings of this study can be accessed through the Sequence Read Archive database. The data is linked to BioProject no. PRJNA1063516 and can be found using accession numbers SRR27547817 to SRR27547879.

## Results

### A titratable tet-inducible CRISPR interference system for *P. aeruginosa*

CRISPR interference (CRISPRi) utilizes dCas9 and sgRNA binding to DNA, forming complexes that effectively repress the expression of the target genes by obstructing transcription. To achieve tunable CRISPRi in *P. aeruginosa*, we developed a tet-inducible system for this bacterium. This system was chosen because of its potential applicability for *in vivo* infection studies. We incorporated a tet-inducible promoter P*_tet_* and a repressor gene *tetR*, codon-optimized for *P. aeruginosa*, into the genetic circuit (**Extended data Fig. 1a**). This configuration allows the transcription of downstream genes driven by P*_tet_* to be regulated by the addition of doxycycline (Dox) known for its tissue penetration ability (Liu et al. 2024). To evaluate the induction efficiency of our tet-inducible system in *P. aeruginosa*, we used the LuxABCDE luminescence reporter system (**Fig. 1a**). Real-time monitoring of P*_tet_*-driven *luxABCDE* expression in PA14 was performed in LB broth with varying concentrations of inducer Dox, a tetracycline derivative. Increasing Dox concentrations (0 to 100 ng/mL), resulted in a dose-dependent elevation in expression levels (**Fig. 1a**), without influence on growth (**Extended data Fig. 1b**). The maximum induction was observed at 100 ng/mL Dox, leading to an approximately 108-fold increase in expression without any discernible growth delay, confirming the efficiency and titratability of the system. We further compared our engineered tet-inducible system to the well-documented arabinose-inducible system in *P. aeruginosa* (McMackin et al. 2021). The tet-inducible system showed a wider dynamic range of induction with less leakage (**Fig. 1a**).

**Fig. 1.**
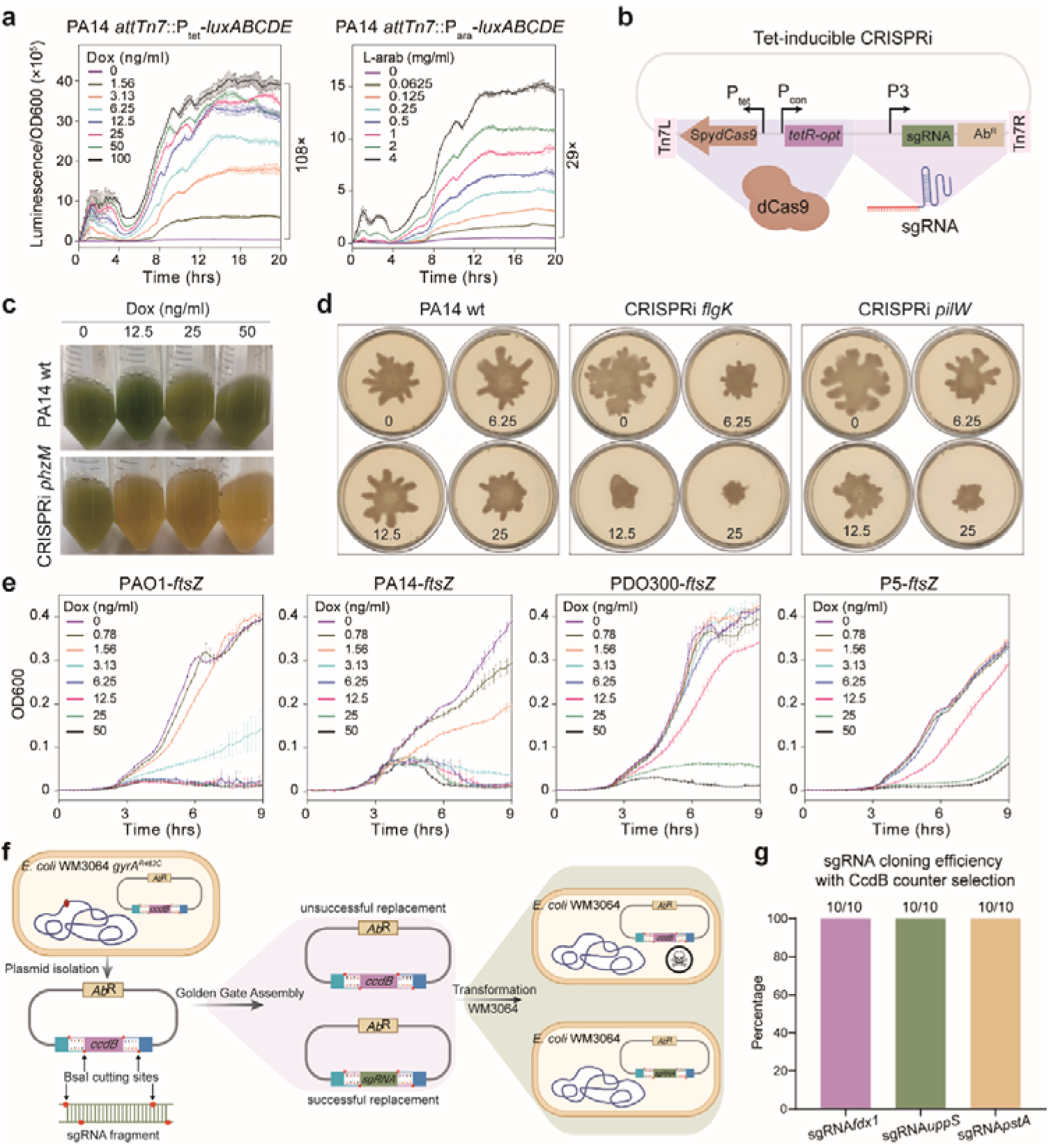
Establishment of tet-inducible CRISPRi system in *P. aeruginosa*. **a.** Evaluation of the tet- and ara-inducible reporter system in *P. aeruginosa*. The efficacy was assessed by induction of *luxABCDE* expression at varying concentrations of doxycycline (Dox) or L-arabinose. **b.** Schematic representation of the tet-inducible CRISPRi system. It employs the SpydCas9-sgRNA complex, which binds to a 20-base pair sequence in the sgRNA-complementary DNA target region, inhibiting transcription elongation via steric hindrance. The *dcas9* is driven by a P*_tet_* promoter, and the sgRNA is expressed under a constitutive promoter. **c.** CRISPRi-mediated repression of pyocyanin biosynthesis. The PhzM is crucial in the phenazine modification pathway and responsible for pyocyanin biosynthesis (a green pigment). **d.** Swarming assay showing the repression of motility by CRISPRi in the presence of Dox (0, 6.25, 12.5, and 25 ng/ml). **e.** CRISPRi represses the expression of the essential gene *ftsZ* in various *P. aeruginosa* strains. **f.** CcdB-based sgRNA cloning strategy. The counter-selection system comprises a *ccdB* gene flanked by BsaI sites for Golden Gate cloning. The toxic protein CcdB acts as a DNA gyrase inhibitor by binding to cleaved double-stranded DNA, causing cell death. Successfully replacing *ccdB* with sgRNAs in WM3064 permits their growth on LB agar plates, significantly enhancing sgRNA cloning efficiency. **g.** Evaluation of the sgRNA cloning efficiency with the CcdB counter selection system. Three sgRNAs were cloned individually,

Next, we employed the tet-inducible system to control the expression of *S. pyogenes dcas9*, to construct a titratable CRISPRi system for *P. aeruginosa* based on the Tn7-based integrative vector pJMP2846 (Banta et al. 2020) (**Fig. 1b**). Previous studies indicated that overexpression of *S. pyogenes* dCas9 is toxic in certain bacteria including *P. aeruginosa* (Cui et al. 2018; Qu et al. 2019). To determine if the expression of dCas9 and integration of the CRISPRi system impacts *P. aeruginosa* growth, we introduced the CRISPRi system into the chromosome of PAO1 and PA14 strains, utilizing an sgRNA targeting *lucff* gene, which is absent in both of the *P. aeruginosa* strains. The outcomes revealed no notable growth impairment across varying degrees of Dox induction (**Extended data Fig. 1c**). We subsequently evaluated the efficiency of the tet-inducible CRISPRi system by targeting several function-known genes, including *phzM*, *flgK*, *pilW,* and *ftsZ,* in *P. aeruginosa*. We turned on the CRISPRi system by addition of Dox to induce expression of dCas9 (CRISPRi ON) and determined its capacity to repress transcription of *phzM*. Gene *phzM* encodes a SAM-dependent methyltransferase, a crucial enzyme in the biosynthetic pathway converting phenazine-1-carboxylic acid to pyocyanin, which imparts the distinctive blue-green pigmentation of *P. aeruginosa* (Wang et al. 2020; Jayaseelan, Ramaswamy, and Dharmaraj 2014). As shown in **Fig. 1c**, CRISPRi-mediated repression of *phzM* transcription resulted in markedly reduced pyocyanin production, with phenotype observed already at a low Dox concentration (12.5 ng/mL). The genes *flgK* (encoding a flagellar hook-associated protein) and *pilW* (encoding minor pilins) were reported to be required for the swarming motility of *P. aeruginosa* (Rattanachak et al. 2022; Marko et al. 2018; Kuchma, Griffin, and O’Toole 2012). We thus assessed the effect of CRISPRi mediated repression of these genes on the cell ability to swarm. Increasing concentrations of Dox resulted in a more significant inhibition of the swarming ability, characterized by smaller flower-like patterns, and reduced number of protrusions (**Fig. 1d**).

To assess the versatility of our CRISPRi system across various *P. aeruginosa* strains, we tested the CRISPRi systems in other strains including reference strain PAO1 (Stover et al. 2000), the mucoid strain PDO300 (Pan, Song, and Ren 2013), and one of our clinical isolates named P5. The CRISPRi was engineered to target the conserved essential gene *ftsZ*, which is crucial for Z ring formation during cell division (Margolin 2005). Repression of *ftsZ* is expected to lead to a growth defect, which was used to evaluate the efficiency of the CRISPRi for an essential gene. As shown in **Fig. 1e**, efficient CRISPRi knockdown of *ftsZ* was achieved in all these strains in a titratable way. However, the concentration of Dox for efficient CRISPRi knockdown differs among the strains, likely due to differences in cell membrane permeability of the inducer. In strains PAO1 and PA14, growth was significantly inhibited with 3.13 ng/mL Dox compared to the no-inducer control (**Fig. 1e**). Conversely, strains PDO300 and P5 required a higher Dox concentration (25 ng/mL) for effective CRISPRi repression. Taken together, these results suggest that the developed tet-inducible CRISPRi system is fully functional and can serve as a foundation for genome-wide CRISPRi libraries in *P. aeruginosa*.

To enable efficient sgRNA cloning for construction of a genome-wide CRISPRi library, we designed a *ccdB* counter selection system in combination with Golden-Gate assembly. This approach facilitates the seamless insertion of the 20 bp base-pairing region of sgRNAs (see Methods) (**Fig. 1f**). The *ccdB* gene, encoding a toxin, was inserted into the cloning site for the sgRNA’s base-pairing region, and the resulting vector was named pCRISPRi-*ccdB*. The pCRISPRi-*ccdB* was maintained in the *E. coli* WM3064 with GyrA462_Arg-Cys_ mutation (SZU148), conferring resistance to CcdB toxicity. The *ccdB* gene fragment was designed with flanking BsaI cleavage sites on both sides, allowing BsaI digestion of the vector to yield ends compatible with the digested sgRNA base-pairing fragments. Following by the Golden Gate assembly, the product was then transformed into *E. coli* WM3064, providing positive selection for vectors in which the sgRNA had successfully replaced the *ccdB* gene. We performed three sgRNA cloning experiments to test the efficiency and achieved a 100% success rate in replacing *ccdB* with sgRNAs (**Fig. 1g**, **Extended Fig. 1d, e**). This indicates that the strategy is effective for generating high-quality sgRNA pools, facilitating construction of compact CRISPRi libraries across *P. aeruginosa* strains.

### Genome-wide CRISPRi-seq in *P. aeruginosa* PA14 refined the essential genes

As demonstrated in **Fig. 2a**, to construct a genome-wide CRISPRi library, we designed sgRNAs for all the annotated genetic features in *P. aeruginosa* PA14 with an automatic sgRNA design pipeline (de Bakker et al. 2022). To improve the robustness of the CRISPRi study, we included two sgRNAs for each target, as the best sgRNAs evaluated by the RNA design pipeline were assigned into set 1, while the 2^nd^ best into set 2. Set 1 consisted of 5981 sgRNAs, whereas set 2 comprised 5971 sgRNAs. The spacers were synthesized as one oligo chip. Two sgRNA pools were cloned in pCRISPRi-*ccdB* vector and maintained in *E. coli* WM3064. PA libraries were constructed by triparental conjugation, introducing the sgRNA libraries into PA14 with the aid of a helper strain harboring the Tn7 transposase plasmid. Both the steps involving sgRNA cloning in *E. coli* WM3064 and the subsequent conjugal transfer from *E. coli* to *P. aeruginosa* were optimized, ensuring the collecting of a number of colonies, resulting in more than 50-fold coverage of sgRNA diversity. Subsequent Illumina sequencing followed by 2FAST2Q analysis (Bravo, Typas, and Veening 2022) indicated establishment of libraries with 5832 sgRNAs in set 1 and 5863 sgRNAs in set 2, covering 97.5% and 98.2% of the genetic features of PA14 strain, respectively (**Supplementary Table S4**).

**Fig. 2.**
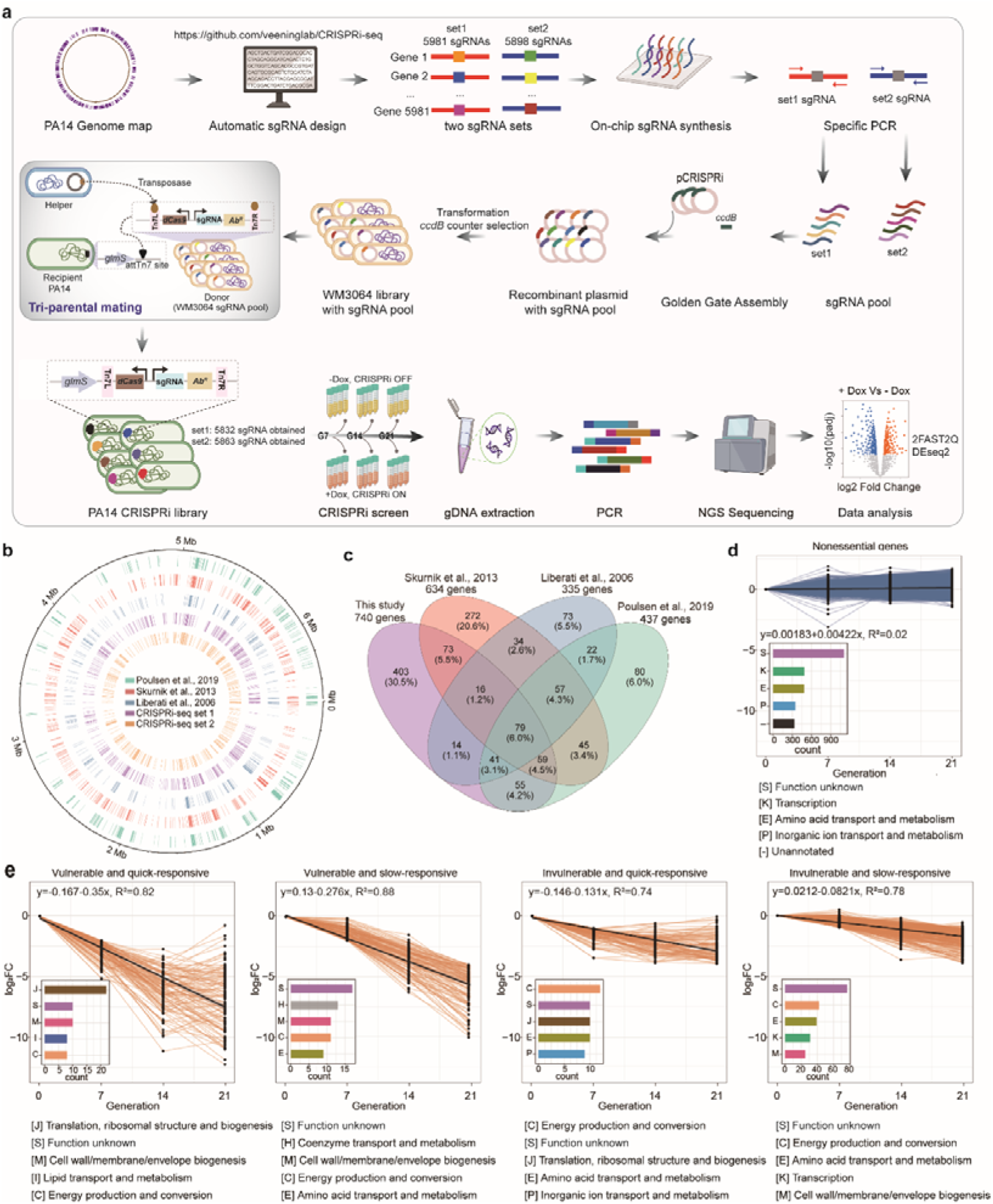
Refinement of essential genes and evaluation of gene vulnerability in *P. aeruginosa* by CRISPRi-seq. **a.** Workflow of the CRISPRi library construction and fitness evaluation. Oligos (set 1 and 2) were synthesized to create sgRNA libraries targeting 5981 and 5898 genes, respectively. These oligos were then amplified by PCR to form double-strand DNAs, which were subsequently cloned into pCRISPRi-*ccdB*. The resulting plasmids were transformed into *E. coli* WM3064 to generate sgRNA libraries, which were integrated in to the PA14 chromosome via triparental mating. Fitness of CRISPRi targets was evaluated at generations 7 (G7), 14 (G14), and 21 (G21) using Illumina sequencing and DEseq2 analysis, with and without CRISPRi activation (Dox +/−). **b.** Genome-wide visualization of CRISPRi-seq outcomes compared to Tn-seq. Log2FC of sgRNAs targeting the PA14 genome is displayed in purple (set 1) and yellow (set 2) in our screen (log2FC < −1). The outer blue, red and cyan tracks represent genes annotated as essential in previous Tn-seq screens reported by Liberati et al. (2006), Skurnik et al. (2013) and Poulsen et al. (2019), respectively. **c.** Comparative analysis of candidate essential genes from CRISPRi-seq and Tn-seq screens. **d, e.** Line plots showing the behavior of sgRNAs targeting the non-essential (**d**) and essential (**e**) genes. Black dots represent sgRNA log2FC values. The solid black line represents the locally estimated scatterplot smoothing fit of the individual mean linear regression at different generations (0, 7, 14 and 21). Bar chart displays Functional classification (top 5) of genes with fitness changes for each cluster, based on available COG terms and manual classifications defined in **Extended data Fig. 2c**.

As a proof of concept, we conducted CRISPRi-seq screenings with both the set 1 and set 2 libraries to refine the list of essential genes in PA14 (**Fig. 2a**). The pooled libraries were cultured in LB broth and maintained in exponential phase by 1:100 back-dilution for 7, 14, and 21 generations with and without addition of Dox for induction. Through a single-step PCR amplification followed by Illumina sequencing, we profiled the abundance of each sgRNA in both induced and non-induced libraries. The fitness quantified by the log_2_FC in sgRNA abundance following induction versus non-induction conditions was then analyzed with the DESeq2 package in R (Love, Huber, and Anders 2014). The sgRNAs exhibiting fitness of log_2_FC less than −1 and adjusted *p*-value under 0.05 were regarded as targeting essential genes. According to this definition, our CRISPRi library screening across 7, 14 and 21 generations revealed 740 candidate essential genes in PA14 strain (**Supplementary Table S5 and S6**). The essential genes delineated by our CRISPRi-seq study exhibit a consistence with those previously identified as essential or responsive through Tn-seq screening methodologies (Liberati et al. 2006; Skurnik et al. 2013; Poulsen et al. 2019) (**Fig. 2b**). However, a clear discrepancy exists among the Tn-seq studies, potentially due to variations in culture conditions or cut-off for essential gene definition. Notably, some essential genes, such as *tRNAs* and the essential housekeeping gene *tuf* encoding translation elongation factor Tu (Whitney et al. 2015), through previous individual mutant studies, were identified exclusively by CRISPRi-seq (**Supplementary Table S6**). This underscores the importance of using CRISPRi-seq to refine the essential genes list in this pathogen. Upon evaluating the relevance of screening results between two CRISPRi libraries over 7, 14, and 21 generations, the correlation coefficient R² remains below 0.05 (**Extended data Fig. 2a**). This is likely due to different degree of transcriptional repression triggered by the two sets of sgRNAs.

Strikingly, the log_2_FC of the majority of sgRNAs targeting essential genes showed a gradual decline with increasing induction generations, indicating that the growth defects caused by the repression of these genes became more pronounced with prolonged CRISPRi repression (**Extended data Fig. 2b**). Interestingly, some essential genes exhibited significant growth defects after just 7 generations of induction, while others only displayed such defects at 14 or 21 generations. This variation suggests that *P. aeruginosa* essential genes respond differently to repression. Based on that, we classified the genes into five categories based on their vulnerability and response time (**Fig. 2d, e**; **Extended data Fig. 2c; Table S6**): 1) nonessential genes, 2) vulnerable and quick-responsive genes, 3) vulnerable and slow-responsive genes, 4) invulnerable and quick-responsive genes, and 5) invulnerable and slow-responsive genes. The representative genes from each category were selected and tested in CRISPRi knockdown growth assays, confirming that the vulnerable and quick-responsive category exhibited the most pronounced and rapid growth inhibition (**Extended data Fig. 3**). COG functional enrichment was performed for the genes classified into each category. Notably, the “translation, ribosomal structure, and biogenesis” and “cell wall/membrane/envelope biogenesis” COG categories were mostly represented in the vulnerable and quick-responsive class. This highlights essential genes in these two categories as the most effective therapeutic targets, consistent with the fact that most used antibiotics target genes in these COGs (Kohanski, Dwyer, and Collins 2010). Taken together, this proof-of-concept study with CRISPRi-seq in PA14 demonstrated the efficiency of this methodology in exploring the biology of *P. aeruginosa*.

### Chemical genetic profiling of gallium in *P. aeruginosa* by CRISPRi-seq

Gallium therapy has been considered as a potential treatment for *P. aeruginosa* infections, but the action mechanisms remain unclear (Goss et al. 2018). To elucidate the genetic factors that modulate gallium’s effectiveness and identify potential targets for enhancing therapeutic efficacy in *P. aeruginosa*, we employed CRISPRi-seq to conduct a genome-wide investigation (**Fig. 3a**). The PA14 CRISPRi library was cultured in LB broth with Dox for 14-generations preinduction before addition of Ga(NO_3_)_3_ for treatment. After addition of Ga(NO_3_)_3_, the bacteria were cultured for another 7 generations of doubling (**Fig. 3a**). The concentration of Ga(NO_3_)_3_ used for the CRISPRi-seq was 200 µM, which leads to 70%-80% of growth reduction (**Fig. 3b**). By using criteria of |log2FC| > 2 and *P*_adj_ < 0.05, we identified 98 sgRNAs whose targeted repression correlated with enhanced fitness, while 9 sgRNAs were associated with reduced fitness, after the treatment with Ga(NO_3_)_3_ (**Fig. 3c**, **Supplementary Table S7**). Gene ontology (GO) enrichment analysis was utilized to discern the distinctive biological characteristics of gallium sensitivity relative genes, which were assorted into various functional categories. There were 98 genes, whose repression caused increased bacterial fitness under gallium treatment, enriched with various translation-related GO terms, such as the large ribosomal subunit, structural constituents of ribosomes, and translation processes (**Extended data Fig. 4a, Supplementary Table S8**). This suggests that the deceleration of protein biosynthesis could potentially mitigate the deleterious effects induced by the gallium treatment. Conversely, the genes whose repression led to reduced fitness were found to be involved in various metabolic processes, including energy metabolism and redox reactions. The enriched ontologies were GTP cyclohydrolase II activity, dihydrolipoyllysine-residue succinyltransferase activity, the oxoglutarate dehydrogenase complex, the P450-containing electron transport chain, ferredoxin-NADP^+^ reductase activity, and the riboflavin biosynthetic process (**Extended data Fig. 4a, Supplementary Table S8**).

**Fig. 3.**
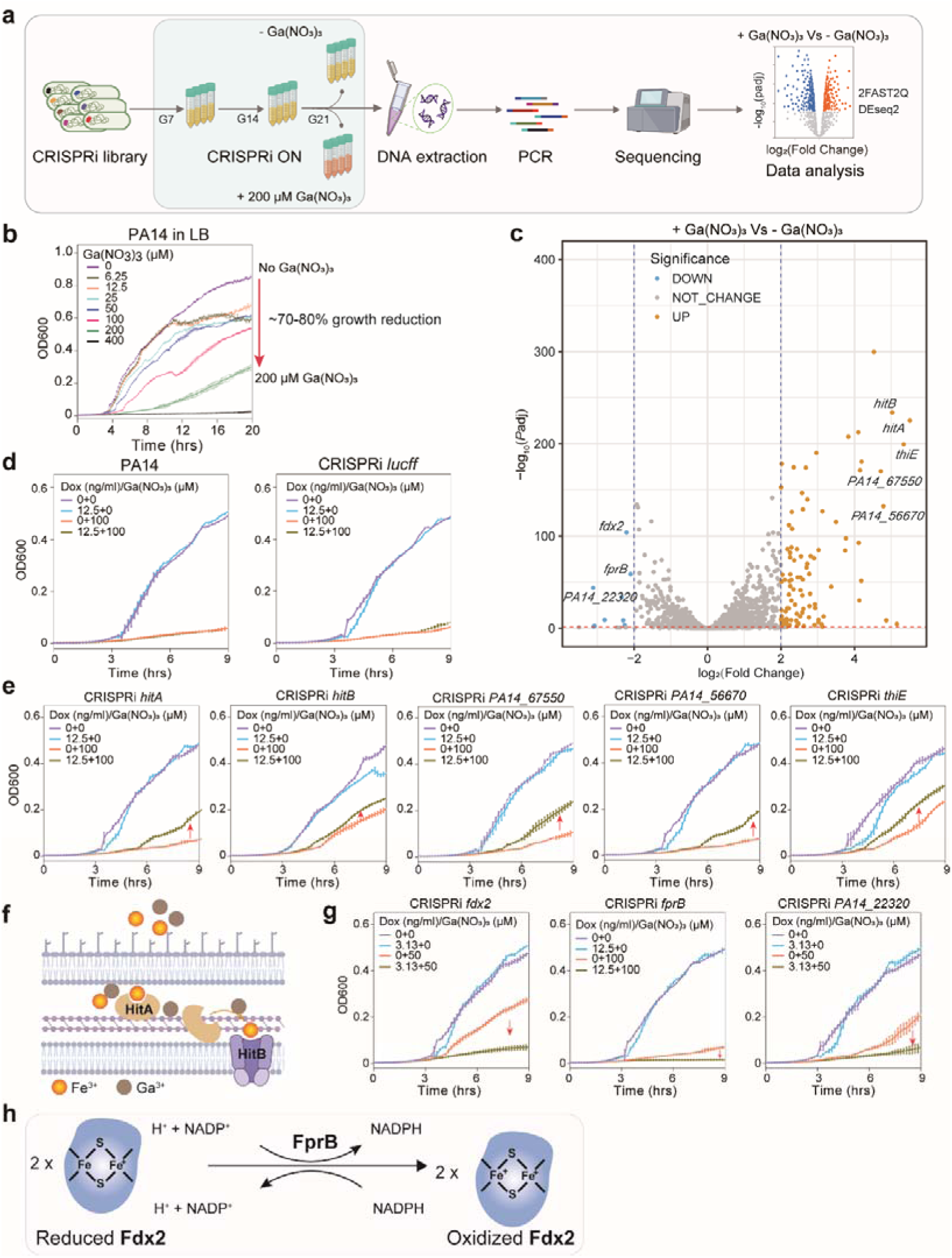
Identification of genes involved in sensitivity to Ga(NO_3_)_3_ therapy using CRISPRi-seq. **a.** Workflow for screening genes involved in sensitivity to Ga(NO_3_)_3_ treatment. The CRISPRi library was induced with Dox (25 ng/ml) until G14, followed by culture with or without 200 μM Ga(NO_3_)_3_ from G14 to G21. **b**. Growth curves of PA14 in LB broth with and without Ga(NO_3_)_3_. **c.** Volcano plots showing the identified genes involved in Ga(NO_3_)_3_ treatment, which plotted gene fitness in log_2_FC and −log_10_(*Padj* value). **d.** Control strains: PA14 and PA14::CRISPRi for non-targeted sgRNA *lucff*. **e.** Validation of down-regulated targets: knockdown strains targeting *PA14_22320*, *fdx2*, *fprB* were constructed with tet-inducible CRISPRi. **f.** Schematic of ABC transporters HitA and HitB involved in Ga^3+^ and Fe^3+^ uptake in *P. aeruginosa*. **g.** Validation of up-regulated group targets: *hitA*, *hitB*, *PA14_67550*, *PA14_56670* and *thiE* were constructed with tet-inducible CRISPRi. **h.** Schematic illustrating the role of FprB in NADP(H)-dependent electron transfer system.

To further explore the molecular mechanisms of *P. aeruginosa*’s response towards gallium treatment, we conducted validation experiments for the top hits identified by CRISPRi-seq. These experiments were performed by assessing the growth of the corresponding knockdown strains in the presence or absence of Ga(NO_3_)_3_. PA14 wild-type and PA14 harboring CRISPRi targeting *lucff* served as controls (**Fig. 3d**). Out of the selected 11 top targets, 8 sgRNAs were successfully validated, showing the reliability of the screening. The gene *hitA*, is top one hit mediating increased fitness under gallium treatment upon repression (**Fig. 3e**), and it encodes a ferric iron-binding periplasmic protein. Notably, our screening also identified *hitB* (**Fig. 3e**), a gene located in the same operon as *hitA*. Consistent with our findings, a previous study identified a loss-of-function mutations in *hitAB* as a major contributor to increased gallium tolerance (Goss et al. 2018). HitAB are involved in Fe^3+^ uptake in *P. aeruginosa*, and were shown to provide the ion binding sites for internalization of gallium in *P. aeruginosa* (**Fig. 3f**) (Guo et al. 2019). Other Fe^3+^ transportation systems were not directly identified in the screening. However, genes whose knockdown increases fitness under gallium stress included *thiE*, in the same operon as *hemL*, required for biosynthesis of heme, as well as *PA14_56670*, encoding an uncharacterized transcriptional regulator and being part of operon encompassing *feoAB* genes, encoding iron transport system. The deletion of *hitA* in PAO1 strain led to drastically increased survival of *P. aeruginosa* upon treatment with gallium (**Extended data Fig. 4b**), indicating that HitAB could be the major transportation system utilized by Ga^3+^ for internalization into *P. aeruginosa* cells.

### The ferredoxin-NADP reductase FprB is crucial for *P. aeruginosa* survival under gallium stress

The genes whose knockdown led to decreased fitness in gallium treatment included *fprB*, encoding a ferredoxin-NADP^+^ reductase; *fdx2,* encoding the ferredoxin 2Fe-2S; and *PA14_22320*, encoding a small hypothetical protein (**Fig. 3g**). Notably, Fdx2 and FprB are involved in the same biochemical reaction in *P. aeruginosa* (**Fig. 3h**). To validate these hits and additionally test the conservation of the observed phenotype in other *P. aeruginosa* strains, we engineered deletion mutants of *PA3237* (ortholog of *PA14_22320*) and *fprB* (*PA4615*) in the PAO1 strain. The gene *fdx2* was identified to be essential in our study and couldn’t be deleted. Although individual CRISPRi strain targeting *PA14_22320* confirmed the results of CRISPRi-seq screening, deletion of this gene did not decrease the tolerance to gallium treatment (**Extended data Fig. 4b**). The deletion of *fprB* did not cause any growth defect in LB agar or LB broth without Ga(NO_3_)_3_ (**Fig. 4a**). However, under the treatment with 5 or 10 μM Ga(NO_3_)_3_, the growth of Δ*fprB* mutant was severely impaired both on LB agar and in LB broth, while the wild-type or complemented Δ*fprB* strain showed no observed growth inhibition with this concentration (**Fig. 4a, b**). Minimum inhibition concentration (MIC) measurements showed that deletion of *fprB* decreased the MIC of gallium from 320 µM to 10 µM (**Extended data Fig. 5a**). Remarkably, the time-killing assay with Ga(NO_3_)_3_ showed that Ga(NO_3_)_3_ acted as a bacteriostatic agent in all the tested concentrations, as high as 1600 µM, towards the wild-type *P. aeruginosa* and complemented Δ*fprB* strain. However, for the Δ*fprB* mutant, as low as 12.5 µM Ga(NO_3_)_3_ led to obvious bactericidal effect (**Fig. 4c**). These results suggest that *fprB* plays an important role in maintaining the survival of *P. aeruginosa* under gallium treatment. Deletion of *fprB* improved the efficacy of gallium treatment towards *P. aeruginosa* not only in terms of inhibition concentration, but also by shifting the killing mode from bacteriostatic to bactericidal. To our knowledge, this study presents the first evidence of an association between *fprB* and gallium susceptibility in *P. aeruginosa*.

**Fig. 4.**
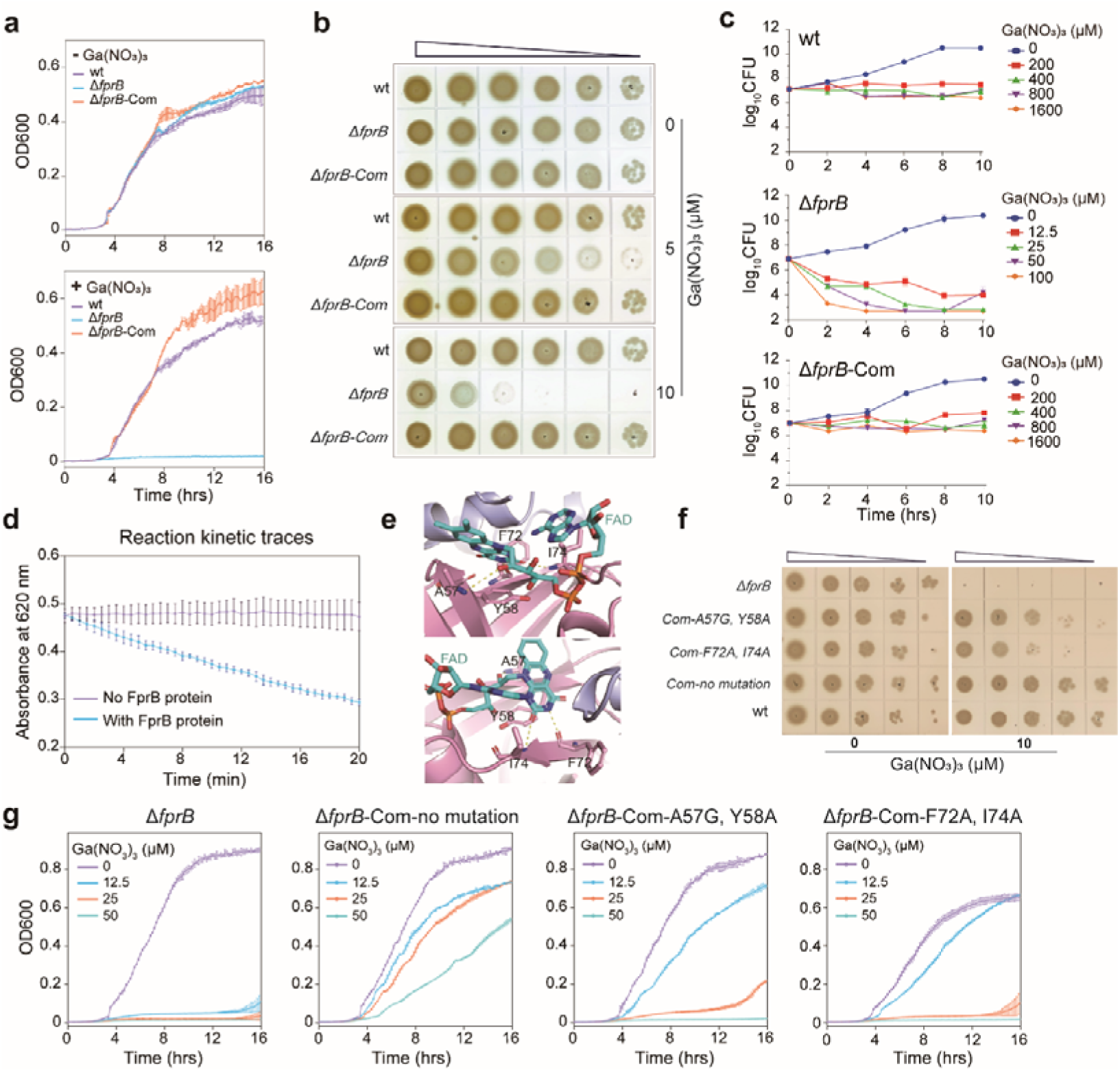
Functional analysis of FprB in Ga(NO_3_)_3_ resistance in *P. aeruginosa*. **a.** Growth curve of PAO1 wild type (wt), *fprB* deletion (△*fprB*) and complementary (△*fprB*-Com) strains in the absence and presence of 12.5 μM Ga(NO_3_)_3_. **b.** Plate spot assays showing sensitivity of PAO1 wt, △*fprB* and △*fprB*-Com strains. **c.** The time-killing assay of by Ga(NO_3_)_3_ towards *P. aeruginosa* in LB broth. Bacterial counts were determined by serial dilutions and plating on LB agar plates. **d.** Reaction kinetic traces for determining FprB activity by 2,6-dichlorophenolindophenol (DCPIP)-diaphorase assay. The experiment was performed at 37□ in a mixture containing 800 nM FprB, 400 μM NADPH, 800 µM DCPIP, 2% DMSO, and 50 mM Tris/HCl buffer (pH 8.0). **e.** Molecular docking of FprB with FAD. Structure of FprB was predicted by Alphafold2 following molecular dynamics refinement. FAD is shown in Corey-Pauling-Koltun (CPK) colored sticks with carbons in light blue. The numbered residues of FprB were predicted near FAD molecule that showed with yellow lines. **f, g.** Impact of FAD binding site mutants (A57G, Y58A, F72A, and I74A) of FprB in *P. aeruginosa*’s resistance to Ga(NO_3_)_3_ treatment. Growth characteristics were determined by tenfold serial dilution followed by dropping onto LB agar plates (**f**) or by monitoring in 96-well plates using a microplate reader (**g**). The initial OD600 value of all strains is 0.003. Data are mean ± SD from triplicate experiments.

The *fprB* encodes a ferredoxin-NADP^+^ reductase (Fpr) that facilitates the reversible electron transfer between NADPH and electron carrier proteins, including [Fe-S] clusters ferredoxin and flavodoxin (Yeom et al. 2009; Romsang et al. 2015; Carrillo and Ceccarelli 2003). To confirm its biochemical activity, we purified the *P. aeruginosa* PAO1 FprB protein and performed the NADPH-dependent 2,6-dichlorophenolindophenol (DCPIP) reduction assay as previously described (Martinez-Julvez et al. 2017). The results showed that FprB efficiently catalyzed the reaction (**Fig. 4d**). Bacterial Fprs contain a prosthetic flavin adenine dinucleotide (FAD), which acts as an electron carrier essential for their catalytic activity (Monchietti et al. 2021; Tondo et al. 2013). Consistently, AlphaFold3 prediction showed that FprB binds to FAD (Abramson et al. 2024) (**Extended data Fig. 5b**). To test whether the observed phenotype of FprB is attributable to its biochemical function, we introduced point mutations into conserved amino acids within the FAD-binding domain of FprB. The mutants: A57G, Y58A, F72A, and I74A, were selected for analysis based on their proximity to the FAD binding site (**Fig. 4e**). As shown in **Fig. 4f, g**, the strains expressing mutated FprB variants exhibited increased sensitivity to gallium compared to both the wild-type and the strain complemented with wild-type FprB in LB agar and LB broth. These findings confirm that the FprB-mediated resistance to gallium therapy depends on its biochemical activity.

### Mechanistic exploration of the increased sensitivity of the **Δ***fprB* mutant to gallium

We next set out to uncover the mechanism of increased sensitivity of *P. aeruginosa* towards gallium caused by inactivation of *fprB*. As the first step, we demonstrated that gallium didn’t inhibit the biochemical activity of FprB by the *in vitro* assay (**Extended data Fig. 5c**). Next, we tested whether deletion of *fprB* leads to altered accumulation of higher concentrations of intracellular gallium. Surprisingly, the *fprB* knockout strains displayed notably reduced intracellular gallium levels in comparison to both wild-type and complemented Δ*fprB* strains (**Extended data Fig. 5d**). Given the antibacterial properties of gallium ions, which interfere with iron-dependent biological processes by attaching to iron-utilizing proteins and competing with iron for bacterial siderophore-mediated uptake (Choi, Britigan, and Narayanasamy 2019), we hypothesized that the deletion of *fprB* may influence the concentration of intracellular iron. Our results demonstrated that in the absence of *fprB* expression, *P. aeruginosa* accumulates higher intracellular Fe and ferrous iron (Fe^2+^) levels compared to wild-type and complemented strains (**Fig. 5a**). The data showed that increased gallium sensitivity of Δ*fprB* was not due to higher intracellular gallium, but rather with severe interference with an iron-dependent biological processes.

**Fig. 5.**
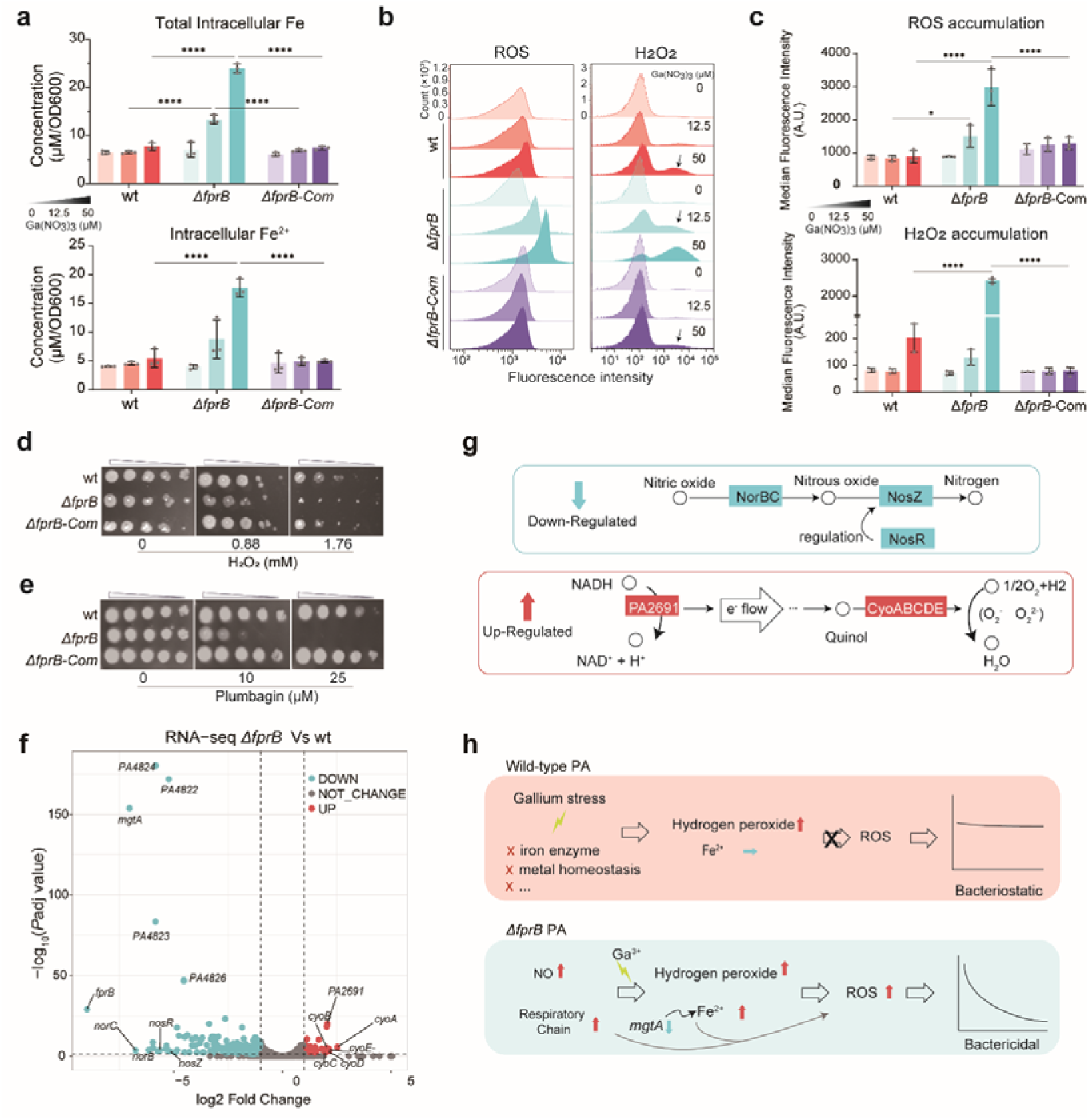
FprB enhances the survival of *P. aeruginosa* by inhibiting Ga(NO_3_)_3_-induced intracellular iron and ROS accumulation. **a.** Intracellular total iron and Fe^2+^ accumulation. Bacteria were cultured for 6 hours in LB broth with or without Ga(NO_3_)_3_, starting at an OD600 of 0.05. Concentration values measured by a microplate reader were normalized to final OD600. **b, c.** ROS and H_2_O_2_ accumulation in bacteria grown for 4 hours in the absence or presence of Ga(NO_3_)_3_, detected by flow cytometry using carboxy-H2DCFDA and Peroxy Orange-1. Data are shown as mean ± SD from triplicate experiments. The asterisks in **a, b** and **c** represent a significant difference from indicated groups with \**P* < 0.05 and \*\*\*\**P* < 0.0001 by two-way ANOVA□and□Tukey’s multiple-comparison test. **d, e.** Plate assays for sensitivity of PAO1 wt, △*fprB* and △*fprB*-Com strains to H_2_O_2_ and Plumbagin. All strains were initially adjusted to an OD600 of 0.003 and then serially diluted tenfold before spotting onto LB agar plates (10^−2^ to 10^−6^). **f.** Volcano plot of differentially expressed genes (|log_2_FC| > 2, *Padj* < 0.05) between △*fprB* and PAO1 wt strains. Up-regulated genes are in red, down-regulated are in cyan, and genes with no significant change in grey. **g.** The role of down-regulated NO reductase (NorBC), N_2_O reductases (NosZ) and NosR, and up-regulated PA2691 and CyoABCDE in bacterial aerobic respiration. **h.** Proposed mechanism of gallium-induced cell death in *P. aeruginosa*. In wild type, gallium can disrupt antioxidant enzymes and metal homeostasis, slightly increasing H□O□ levels and resulting in a bacteriostatic effect. In *fprB*-deficient *P. aeruginosa*, heightened NO levels and aerobic respiration, coupled with the increased intracellular iron, trigger the Fenton reaction, resulting in rapid ROS accumulation and bactericidal effects.

Increased intracellular ferrous ions can react with hydrogen peroxide via the Fenton reaction, leading to the accumulation of reactive oxygen species (ROS), which may cause severe oxidative damage and cell death (Belenky et al. 2015; Vilchèze et al. 2013). To test whether the increased gallium sensitivity of the Δ*fprB* mutant is due to increased ROS, we measured intracellular hydrogen peroxide using Peroxy Orange-1 and ROS using carboxy-H2DCFDA, the specific fluorescent indicator dyes (**Fig. 5b, c**). The analysis showed that gallium treatment induced hydrogen peroxide production in both wild-type and Δ*fprB* mutant strains, with higher production in Δ*fprB* mutant. Specifically, the wild-type strain treated with 50 µM Ga(NO_3_)_3_ produced similar levels of hydrogen peroxide as the Δ*fprB* mutant treated with 12.5 µM Ga(NO_3_)_3_. No detectable hydrogen peroxide was observed in the *fprB*-complemented strains, possibly due to FprB overexpression. These findings indicate that FprB plays a role in controlling hydrogen peroxide production under gallium treatment. Furthermore, ROS accumulation was observed exclusively in the Δ*fprB* mutant, but not in the wild-type or complemented strains, under the tested gallium concentrations. This finding underscores the role of FprB in regulating ROS levels and its contribution to gallium sensitivity. Consistently, intracellular concentration of NADP^+^ and NADPH was significantly increased in the Δ*fprB* mutant under gallium treatment compared to wild-type or complemented strain (**Extended data Fig. 6a**), indicating an increase of oxidative stress in the knockout strain (Singh et al. 2007). The elevated NADP^+^/NADPH ratio in the Δ*fprB* mutant under gallium stress further demonstrated the loss of a reduced redox state in the mutant (**Extended data Fig. 6a**). Notably, supplementing NADPH to the medium increased the survival of the Δ*fprB* mutant upon gallium exposure (**Extended data Fig. 6b**). Further confirming the role of FprB in controlling ROS, we found that Δ*fprB* mutant showed increased sensitivity to hydrogen peroxide and plumbagin, which is a stimulator of ROS generation (Hong et al. 2019) (**Fig. 5d, e**).

To further elucidate the mechanism by which FprB controls ROS production, we performed RNA-seq analysis (**Fig. 5f** and **Supplementary Table S9**). The analysis revealed that several genes related to nitric oxide reduction were significantly down-regulated, such as *norB*, *norC*, *nosZ*, and *nosR*, in the Δ*fprB* mutant. NorBC catalyzes the two-electron reduction of NO to N_2_O, while NosZ is the only enzyme known to reduce N_2_O to N_2_ with NosR as the electron donor for N_2_O reduction (**Fig. 5g, upper panel**). Inhibition of the denitrification respiratory pathway can elevate NO levels, a reactive nitrogen species (RNS) free radical, leading to nitrosative stress and subsequent cell injury and death (Corpas, Río, and Palma 2019; Zhang et al. 2019). In addition, *mgtA* (*PA4825*) and the neighboring genes which might compose the same operon (*PA4822-24*) were also found to be significantly downregulated in the Δ*fprB* mutant. It was reported that loss function of *mgtA* leads to accumulation of intracellular Fe (Marshall et al. 2009), which explains the observation of **Fig. 5a**. The genes up-regulated in the Δ*fprB* mutant includes *PA2691*, *cyoA*, *cyoB*, *cyoC*, *cyoD*, and *cyoE*. PA2691 is an NADH dehydrogenase that catalyzes electron transfer from NADH to quinone, serving as a key entry point for electrons in the respiratory chain (Kordes et al. 2019). The cytochrome o complex (CyoABCDE) is an ubiquinol oxidase in bacterial aerobic respiration. It catalyzes the two-electron oxidation of ubiquinol-8 and the four-electron reduction of O_2_ to H_2_O, coupling electron flux to proton motive force generation across the membrane (Salunkhe et al. 2005) (**Fig. 5g, lower panel**). Based on the above information, we proposed a model to elucidate the increased gallium sensitivity and bactericidal effect observed in the Δ*fprB* mutant (**Fig. 5h**): Upon uptake by *P. aeruginosa*, gallium disrupts iron enzymes and metal homeostasis, triggering hydrogen peroxide production. In the wild-type strain, detoxification enzymes degrade hydrogen peroxide, preventing further ROS formation and subsequent cell death. However, in the Δ*fprB* mutant, gallium-induced hydrogen peroxide reacts with increased intracellular iron, fueling the Fenton reaction and leading to accumulation of ROS. Concurrently, the mutant exhibits a higher basal level of nitric oxide due to downregulated *nor* genes and an enhanced respiratory chain from upregulated electron transfer genes, potentially exacerbating ROS leakage. The accumulation of ROS likely hastens cell death, thereby explaining the bactericidal action of gallium against the Δ*fprB* mutant.

Notably, there is another ferredoxin-NADP^+^ reductase encoding gene *fprA*, which is an essential gene (**Extended Data Fig. 7d**), in the chromosome of *P. aeruginosa*, encoding a protein which shares 42% amino acid sequence identity with FprB in both PAO1 and PA14 strains (**Extended Data Fig. 7f, g**). However, neither FprA nor its substrate Fdx1 was identified in our CRISPRi-seq screen under gallium stress (**Extended Data Fig. 7d**). Individual *fprA* repression mutant did not significantly increase sensitivity to gallium (**Extended Data Fig. 7e**), and complementing *fprA* in the Δ*fprB* mutant did not enhance gallium tolerance (**Extended Data Fig. 7h**). These findings suggest that although FprA and FprB are both ferredoxin-NADP^+^ reductases, they function in distinct pathways.

### The ferredoxin-NADP^+^ reductase FprB is a promising therapeutic target for *P. aeruginosa* infection

Given the hypersensitivity of the *P. aeruginosa* Δ*fprB* mutant to gallium therapy in vitro, we investigated FprB as a potential synergistic target for gallium therapy. A comprehensive genome analysis of all the 981 *P. aeruginosa* strains with full genomes on NCBI revealed that all but one contained the *fprB* gene ortholog, with each strain harboring a single copy (**Fig. 6a**). The genomic context of *fprB*, including the order and transcription directions of flanking genes, was consistent across different genomes (**Fig. 6b**). The protein sequences of FprB showed minimal variance (**Extended Data Fig. 7a**). Despite slight variations, the genes exhibited significant collinearity (**Extended Data Fig. 7b**), indicating a unique evolutionary origin without horizontal transfer. The FprB protein sequences were highly conserved, with only 20 polymorphic sites detected among the 258 amino acid positions in representative strains from various phylogenetic sub-clusters (**Extended Data Fig. 7c**). The largest variation still maintained 95.7% identity with the consensus sequence. These findings confirm that FprB is highly conserved in *P. aeruginosa* and can be targeted in most strains.

**Fig. 6.**
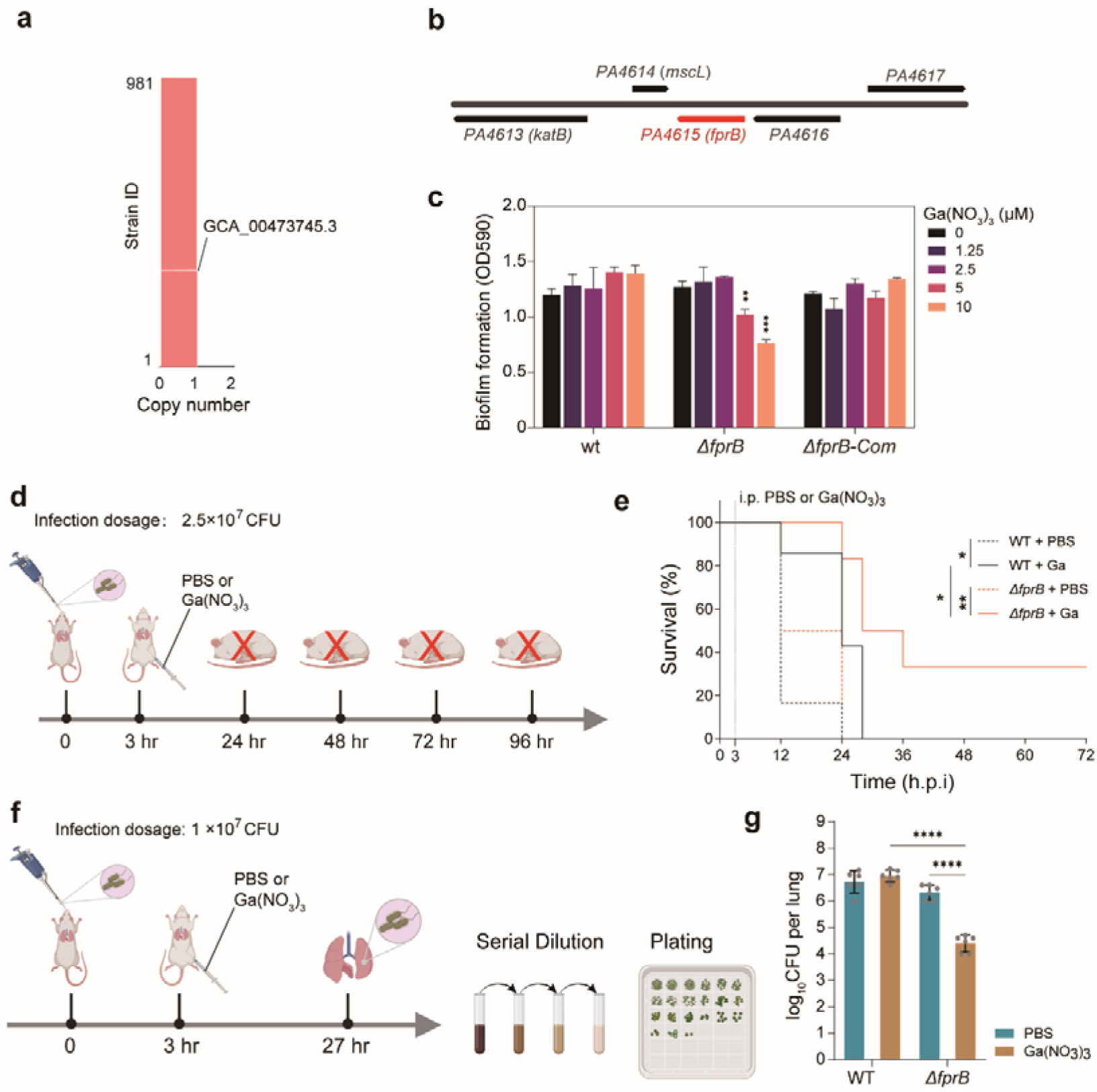
FprB is a promising synergistic target of gallium therapy in *P. aeruginosa*. **a.** Genome analysis of 981 *P. aeruginosa* strains from NCBI shows the presence of the *fprB* gene in most strains except GCA_00473745.3 with a single copy. **b.** Genomic context analysis of the *fprB* gene in 981 *P. aeruginosa* strains reveals consistent gene order across different genomes. **c.** Biofilm formation by *P. aeruginosa* with or without Ga(NO□)□ assessed using the crystal violet method after for 24-hour incubation at 37°C. **d, e.** Survival of lung infected mice. BALB/c mice were intranasally administrated with *P. aeruginosa* (2.5 × 10^7^ c.f.u.). At 3 h.p.i., mice were treated with 50 µL µL of PBS or 250 mM Ga(NO□)□ via intraperitoneal injections. **f, g.** Bacterial load in lung tissues of infected mice. BALB/c mice (female, 6-8 weeks) were intranasally administrated with *P. aeruginosa* (1.0 × 10^7^ c.f.u.). At 3 h.p.i., mice were treated with 50 µL of PBS or 250 mM Ga(NO□)□ via intraperitoneal injections. At 27 h.p.i., mice were sacrificed to assess the bacteria load in the lung. The asterisks in **c** and **g** represent a significant difference from indicated groups with \**P* < 0.05, \*\**P* < 0.01, \*\*\**P* < 0.001 and \*\*\*\**P* < 0.0001 by Gehan-Breslow-Wilcoxon test (**e**) or two-way ANOVA□and□Tukey’s multiple-comparison test (**c**, **g**).

*P. aeruginosa* is known to form biofilms in CF airways (Bjarnsholt et al. 2009). We tested the efficiency of gallium in inhibiting biofilm formation in both wild-type and Δ*fprB P. aeruginosa* strains. Gallium effectively inhibited biofilm formation in the Δ*fprB* mutant at concentrations as low as 5 to 10 µM, which is close to the peak plasma and sputum concentrations in humans (Goss et al. 2018). In contrast, the wild-type and complemented strain were not similarly affected (**Fig. 6c**). To further evaluate FprB as a synergistic target for gallium therapy in vivo, we utilized a murine lung infection model (**Fig. 6d, f**). Deletion of *fprB* in *P. aeruginosa* significantly increased animal survival and reduced bacterial burden in the lungs under gallium therapy (**Fig. 6e, g**). These results highlight the great potential of targeting FprB to enhance the therapeutic efficacy of gallium against *P. aeruginosa* infections.

## Discussion

Our study establishes and validates a tet-inducible pooled CRISPRi-seq screening strategy to uncover the genetic basis of growth-related traits in the human pathogen *P. aeruginosa*. By applying CRISPRi-seq to profile the chemical genetics of gallium treatment, we identified genes associated with gallium susceptibility on a genome-wide level. Notably, we identified FprB as a synergistic target for gallium therapy, demonstrating the effectiveness of CRISPRi-seq based genome-wide forward genetic screenings. Future research should focus on developing drug-like compounds targeting FprB to improve the selectivity and efficacy of gallium-based treatments.

The CRISPRi technique has increasingly become a popular tool for exploring gene functions in various bacteria under given conditions (Qi et al. 2013; Bikard et al. 2013; Liu et al. 2017; Wang et al. 2018; Lee et al. 2019). However, its application in *P. aeruginosa* is rare, and no genome-wide scale study utilizing CRISPRi-seq was described. In contrast, several genome-wide analyses of gene function in *P. aeruginosa* employing Tn-seq techniques have been reported (Gallagher, Shendure, and Manoil 2011; Lee et al. 2015; Skurnik et al. 2013; Poulsen, Clatworthy, and Hung 2022). The transposon insertion techniques have certain limitations, such as the lack of targeted insertion capability, applicability restricted to non-essential genes, and a tendency for polar effects (Liu et al. 2021; Zhu et al. 2023; de Bakker et al. 2022). The engineered tet-inducible CRISPRi system presented in this study could provide robust reinforcement for the transposon-based techniques. Our genome-wide CRISPRi-seq study in *P. aeruginosa* PA14 successfully refined the list of essential genes in this strain. By designing sgRNAs for all annotated genetic features and constructing compact CRISPRi libraries, we achieved high coverage and robustness in our study. The inclusion of two sgRNAs per target gene allowed for a more thorough investigation, and our optimized cloning and conjugation processes ensured a diverse and representative library. Through CRISPRi-seq screenings, we identified 740 candidate essential genes, many of which align with previously identified essential genes from Tn-seq screenings in the same strain (Liberati et al. 2006; Skurnik et al. 2013; Poulsen et al. 2019) (**Fig. 2b, c**). Notably, some essential genes were uniquely identified through CRISPRi-seq screening, highlighting the distinct advantages and necessity of this approach. However, the potential for polar effects of the CRISPRi system on genes within the same transcriptional unit (Cui et al. 2018; Qi et al. 2013), which may lead to false positives, must be considered. To address this, we incorporated operon information for each gene in *P. aeruginosa* PA14 (Supplementary Table S6), enabling a more accurate interpretation of the CRISPRi-seq data. The quantitative and time-dependent fitness measurements allowed us to classify essential genes based on their vulnerability to CRISPRi repression. Such categorization of essential genes offers valuable insights into their potential as therapeutic targets (Hawkins et al. 2020; Bosch et al. 2021) (**Fig. 3d, e**). Notably, genes from COGs of “translation, ribosomal structure, and biogenesis” and “cell wall/membrane/envelope biogenesis” were identified as the mostly represented COGs in the “vulnerable and quick-responsive” category. These findings highlight these genes as the most promising targets for antimicrobial development. In fact, the majority of the clinically used antibiotics target genes within these two COG categories (Kohanski, Dwyer, and Collins 2010). Additionally, 27 genes categorized as “vulnerable” fall under the “Function unknown” COG category (**Supplementary Table 6**). Further research is needed to elucidate the functions of these 27 genes or the operons they belong to, as they may represent key targets for *P. aeruginosa* therapy. In summary, this proof-of-concept study highlights the efficacy of CRISPRi-seq in dissecting the genetic landscape of *P. aeruginosa*, paving the way for novel therapeutic strategies. Future studies should focus on refining the gene vulnerabilities under different growth conditions, including those which are relevant *in vivo*.

In this study we focused on chemical genetic profiling via CRISPRi-seq to elucidate genetic factors that influence the efficacy of gallium therapy against *P. aeruginosa*. We identified genes whose repression either enhanced or diminished bacterial fitness under gallium treatment. Key findings include the identification of *hitA* and *hitB*, both involved in Fe^3+^ uptake, which confer increased fitness when repressed, corroborating earlier findings in *P. aeruginosa* (Guo et al. 2019; Zemke et al. 2020). The identification of ferric ion-associated mechanisms is not surprising, as gallium and ferric ion share comparable radii and chemical properties, allowing gallium ion to be incorporated via the route of the *P. aeruginosa* iron uptake system involving HitA and HitB and to act as an iron mimic in the biological processes (Zemke et al. 2020; García-Contreras et al. 2013). The identification of HitAB, but not other iron uptake systems, indicates HitAB is the major transportation system utilized by gallium.

Our work represents the initial studies to utilize CRISPRi-seq to identify genes whose loss of function results in increased sensitivity to gallium. Notably, the deletion of *fprB* led to a substantial reduction in the minimum inhibitory concentration (MIC) of gallium, from 320 µM to 10 µM. This reduced MIC falls within the range of peak plasma and sputum concentrations observed in humans (8 ~ 12 µM) (Goss et al. 2018), underscoring the potential of *fprB* as a promising therapeutic target. It has been demonstrated that gallium can prompt an increased intracellular ROS level (Li et al. 2021; Guo et al. 2023). The gene *fprB* encode a ferredoxin-NADP^+^ reductase (FprB). FprB binds FAD and NADP(H) within its N- and C-terminal domains, playing a crucial role in detoxifying ROS and maintaining redox homeostasis (Monchietti et al. 2021; Carrillo and Ceccarelli 2003). Mechanism exploration showed the Δ*fprB* mutant displayed elevated intracellular iron and ferrous iron (Fe^2+^) levels (**Fig. 5a**), which might react with the gallium induced ROS, especially hydrogen peroxide, and lead to production of more dangerous radical ROS via the Fenton reaction (Vilchèze et al. 2013; Belenky et al. 2015). This is in consistent with the observation of higher hydrogen peroxide and ROS levels in the Δ*fprB* mutant under gallium treatment, compared to wild-type and complemented strains (**Fig. 5b, c**). RNA-seq analysis of the transcriptomes of the Δ*fprB* and wild-type strains revealed upregulation of genes associated with aerobic respiration and downregulation of those involved in nitric oxide reduction in the Δ*fprB* mutant. These transcriptional changes likely exacerbate ROS production, further contributing to cell death. Previous study showed that gallium exhibits bacteriostatic mode of action against *P. aeruginosa* by affecting the key enzymes in multiple metabolic pathways (Wang et al. 2023). Remarkably, our findings revealed a shift in gallium’s mode of action from bacteriostatic to bactericidal upon the loss of the *fprB* gene. This highlights how the genetic background of bacteria can significantly influence the effectiveness of an antimicrobial, transforming its impact from merely inhibiting growth to actively killing the pathogen. These insights offer a promising strategy to enhance gallium therapy by exploiting specific genetic vulnerabilities in *P. aeruginosa*. Lastly, considering that gallium therapy for cystic fibrosis (CF) patients suffering from chronic *P. aeruginosa* infections is currently in Phase II clinical trials and has shown promising outcomes, we explored the potential of FprB as a synergistic target to enhance the efficacy of this treatment with CF infection related models. Given that *P. aeruginosa* forms resilient biofilms in the CF airway, which are a major obstacle to effective treatment (Bjarnsholt et al. 2009), we performed biofilm inhibition assays and utilized a murine lung infection model to evaluate the impact of targeting FprB. Our results demonstrated that *P. aeruginosa* strains lacking *fprB* showed significantly reduced biofilm formation, even at low gallium concentrations (5 ~ 10 μM). Furthermore, in the murine lung infection model, the absence of *fprB* significantly enhanced the therapeutic efficacy of gallium. Together, this study provides deeper insight into the genetic interactions with gallium, paving the way for more effective therapeutic strategies against *P. aeruginosa* infections.

## Contributions

Y.Z. and X.L. designed the research. Y.Z., T.T.Z., X.X., J.Z.O., A.M.R., W.H.S., Y.J.L., and Y.L.L. performed the experiments and analyzed the data. A.K., V.D.B., L.Y., L.Y., N.J., J.W.V., Y.J.W., and X.L. contributed to data analysis. Y.Z. and T.T.Z. analyzed the sequencing data. Y.J.W. performed the evolutionary conservation analysis. Y.Z. and X.L. wrote the manuscript. X.L. supervised the project. All authors reviewed and edited the manuscript.

## Acknowledgments

We thank Prof. Sisi Li, Dr. Xuan Du, and Dr. Fan Zou from Shenzhen University for helpful discussions and technical support, Dr. Xueling Lu from University of Groningen for help in data analysis. Furthermore, we thank Prof. Xilin Zhao from Xiamen University for insightful discussions. This work was supported by the National Key Research and Development Program of China (2023YFD 10800100); National Natural Science Foundation of China (82200047, 82270012); the Science and Technology Project of Shenzhen (JCYJ20220818095602006); Shenzhen Science and Technology Program (KQTD20200909113758004); Pearl River Talent Project of Guangdong Province (2021QN02Y283); Shenzhen University 2035 Program for Excellent Research (86901-00000216).

## Extended Data

### Extended Figures

**Extended data Fig. 1.**
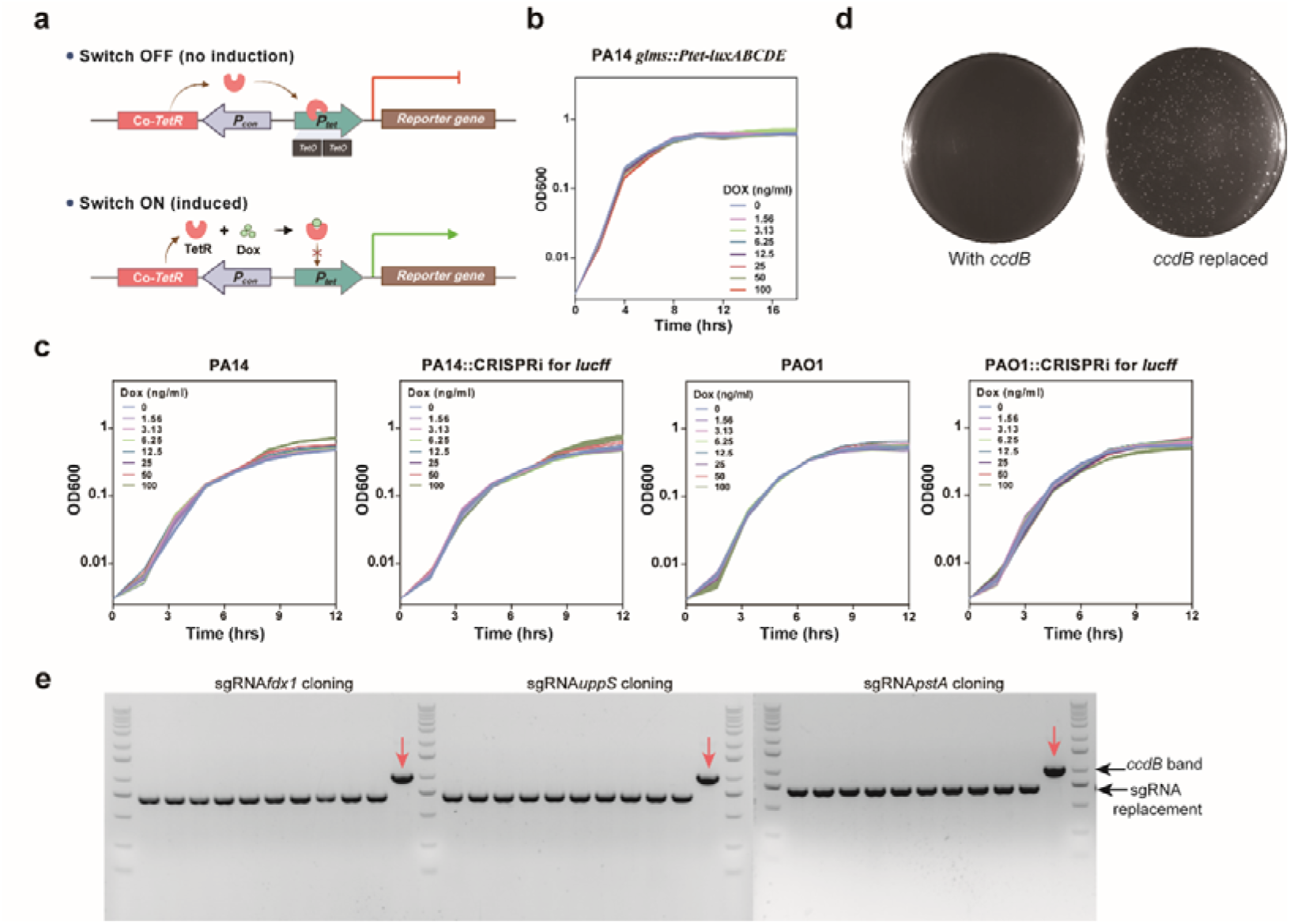
Evaluation of the tet-inducible CRISPRi system and CcdB-counter selection for sgRNA cloning. **a.** The designed tet-inducible system for P. aeruginosa. It includes a constitutive promoter (P*_con_*) driving a codon-optimized TetR and a tet-inducible promoter (P*_tet_*) initiating reporter gene expression. Dox presence inhibits TetR binding to *tetO* on P*_tet_*, enabling reporter gene transcription. **b.** Impact of doxycycline on growth of *P. aeruginosa* PA14 with tet-inducible system. **c.** Impact of CRISPRi system on growth of *P. aeruginosa*. The *P. aeruginosa* strains PAO1 and PA14 were engineered to express a CRISPRi system including dCas9 to repress transcription of the luciferase reporter gene *lucff*. As *lucff* is absent in the PAO1 and PA14 genomes, this allows quantification of any growth defects resulting specifically from dcas9 expression. **d, e.** Enhanced sgRNA cloning efficiency via *ccdB* replacement in bacteria. The successful replacement of *ccdB* by sgRNAs in bacteria allows for their survival on LB agar plates (**d**). 100% positive sgRNA cloning efficiency was achieved as validated by colony PCR (**e**).

**Extended data Fig. 2.**
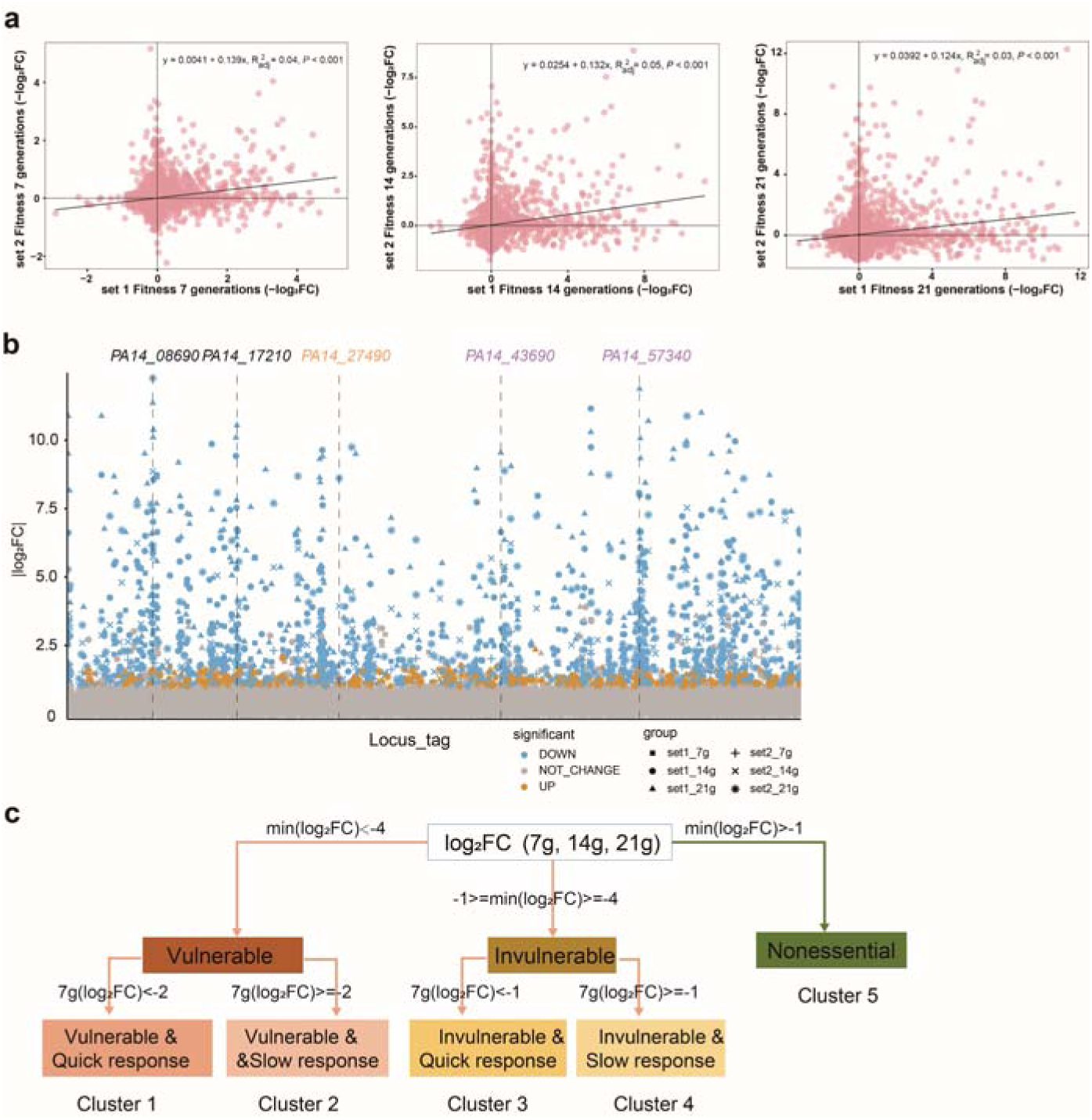
Assessment of the gene essentiality in *P. aeruginosa* via CRISPRi-seq. **a.** Correlation of gene fitness evaluated by set 1 and set 2 libraries, across 7, 14 and 21 generations of Dox induction. **b.** Comparative analysis of essential gene targets from set1 and set 2 libraries. Locus-tags in black (e.g. *PA14_08690* and *PA14_17210*) were screened in both libraries. Those in purple (*PA14_43690* and *PA14_57340*) were unique to set 1 library, while the one in yellow (*PA14_27490*) was only screened by set 2 library. **c.** The workflow for categorizing genes into five clusters based on response time and maximum log_2_FC in PA14 utilizing CRISPRi-seq.

**Extended data Fig. 3.**
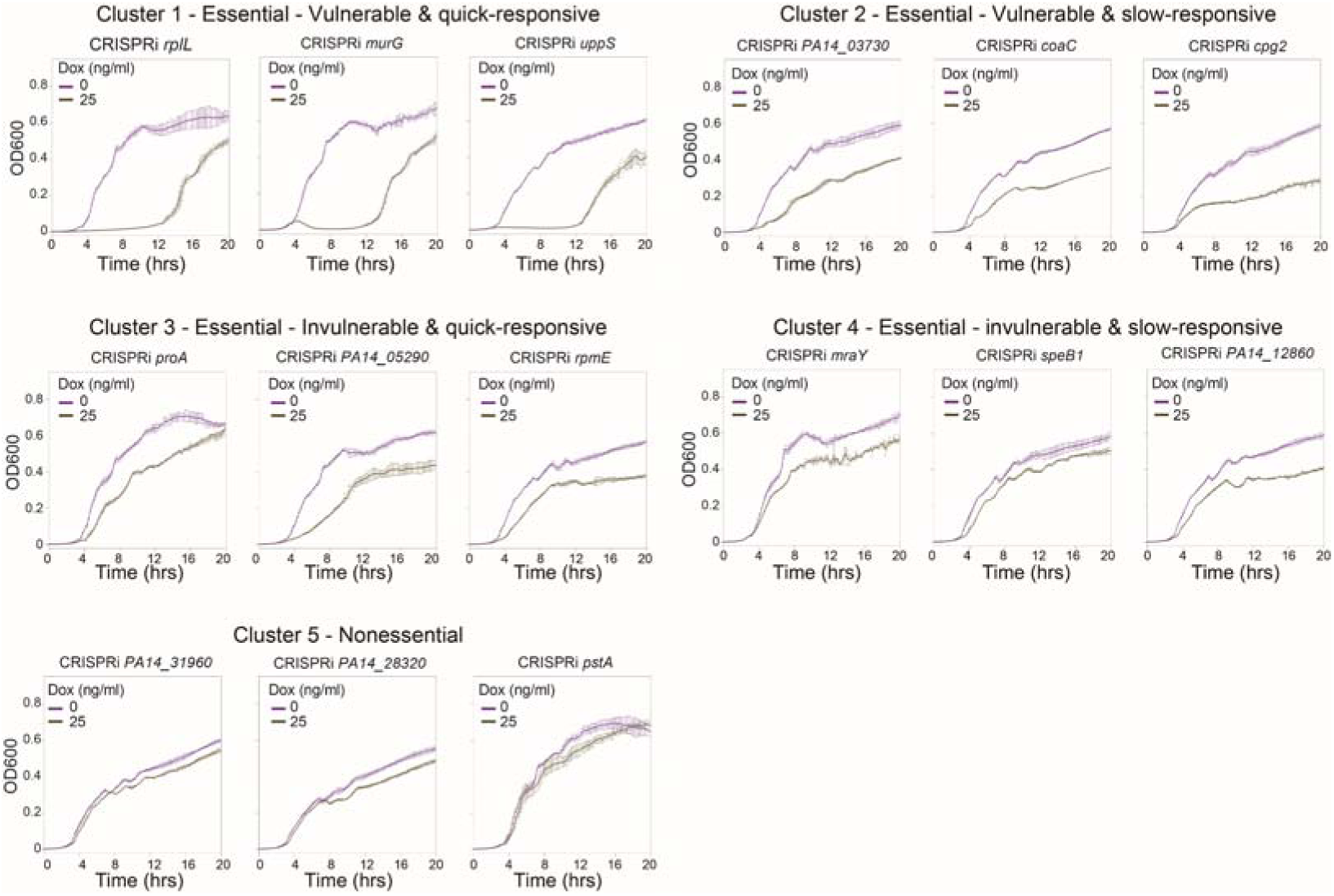
Validation of growth for genes from the 5 clusters identified in Extended data Fig. 2c. All strains were initially adjusted to an OD600 of 0.003. Experiments were performed in triplicate and data are presented as mean ± SD.

**Extended data Fig. 4.**
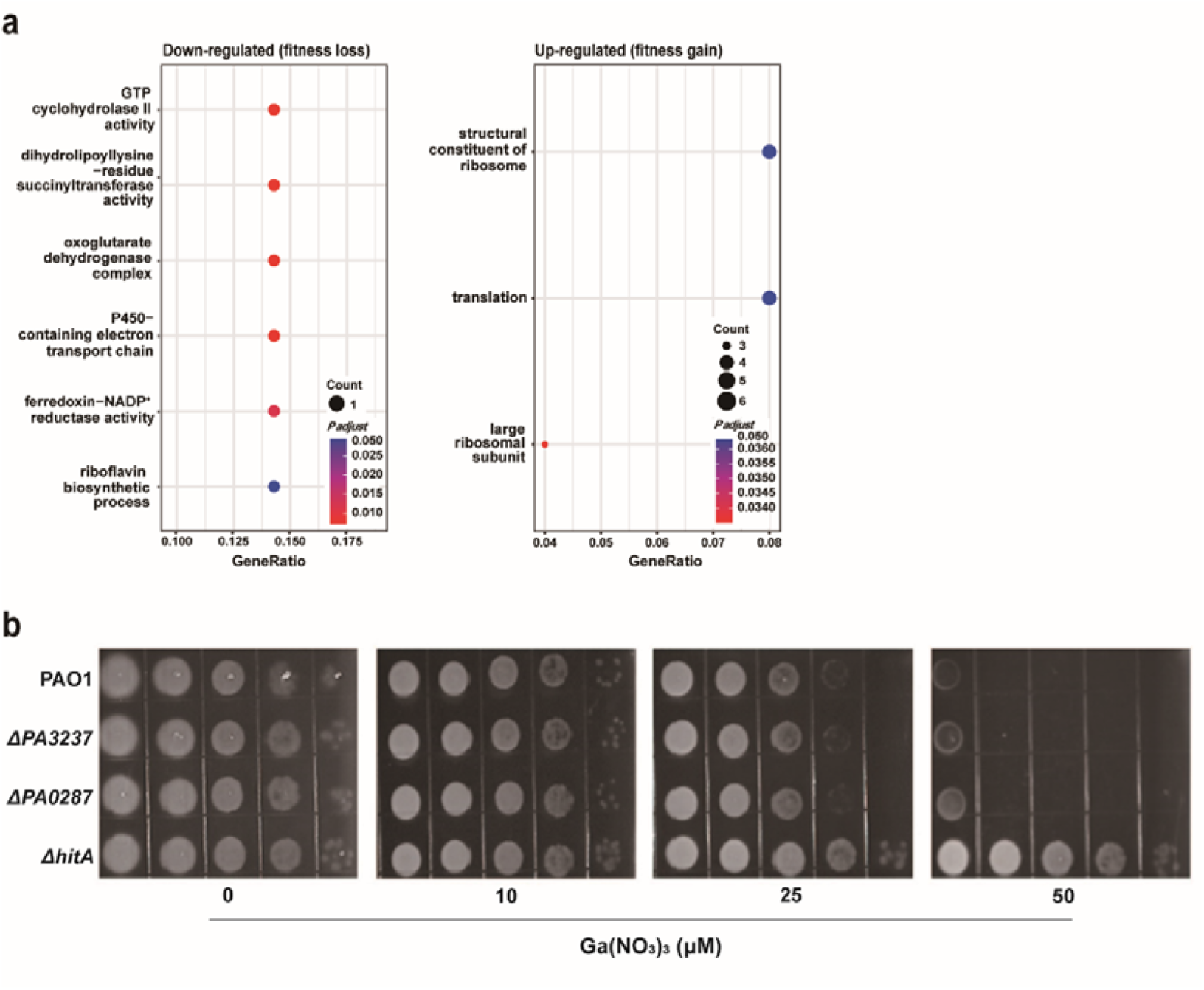
Impact of Ga(NO_3_)_3_ on *P. aeruginosa* growth. **a.** GO enrichment analysis of the genes with differential fitness in Ga(NO_3_)_3_-treated and untreated *P. aeruginosa,* identified by CRISPRi-seq. **b.** Deletion of *PA3237* (ortholog of *PA14_22320*) and *PA0287* (ortholog of *PA14_03760*) shows no growth difference under gallium treatment compared to wild-type PAO1. In contrast, deletion of *hitA* increases PAO1 resistance to gallium.

**Extended data Fig. 5.**
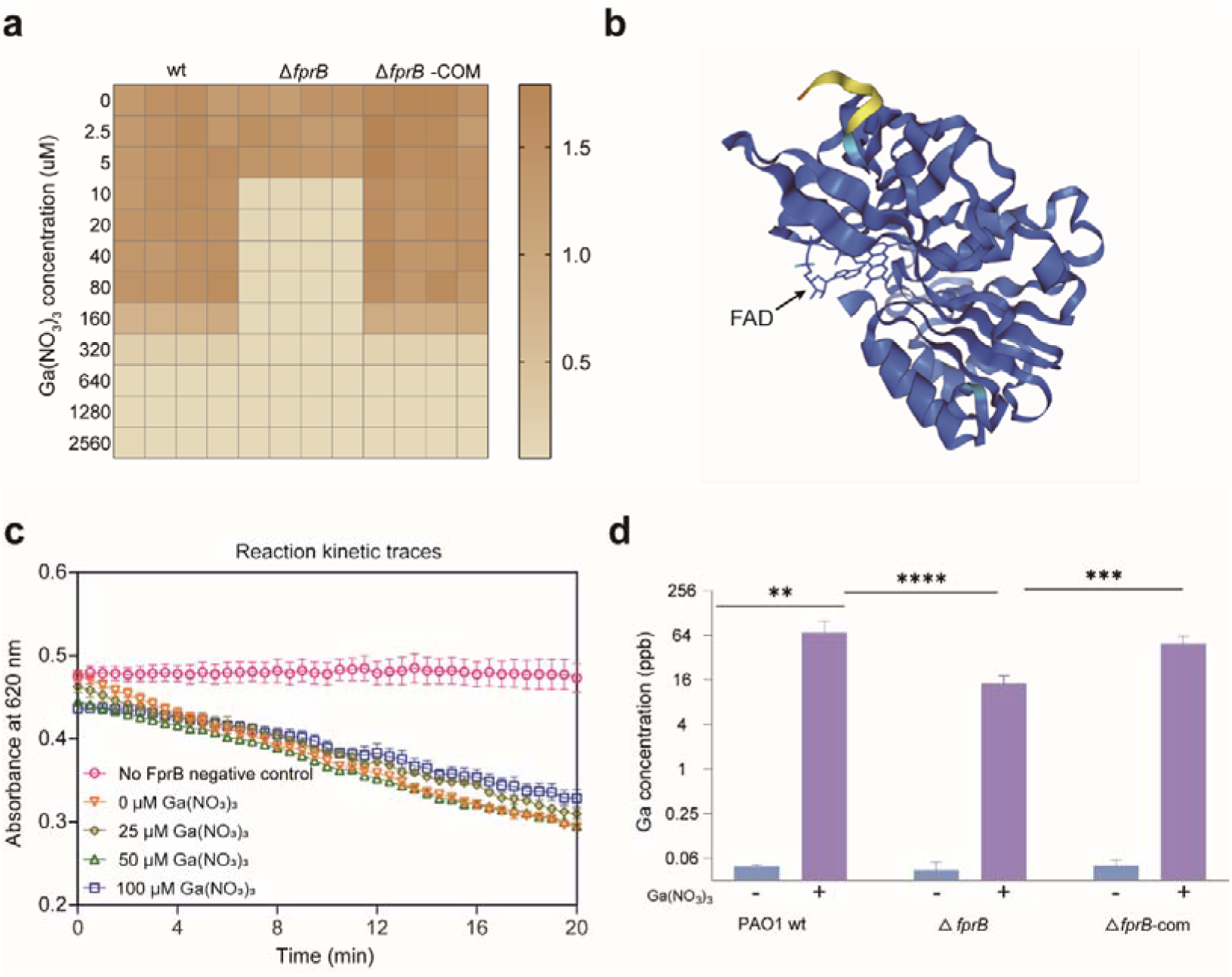
Functional analysis of FprB. **a.** MIC determination of Ga(NO_3_)_3_ for *P. aeruginosa.* Bacteria were cultured in 96-well plates for 24 hours at 37°C with 4 replicates per strain. **b.** AlphaFold3 prediction showing FprB binding to FAD. **c.** Dose-response curves for the FprB 2,6-dichlorophenolindophenol (DCPIP)-diaphorase activity in the absence or presence of Ga(NO_3_)_3_. Experiments performed at 25°C in a mixture containing 800 nM FprB, 800 µM DCPIP and 400 µM NADPH in 50 mM Tris/HCl pH 8.0 (n = 3, mean ± SD). Negative control contains all reactants but no FprB. **d.** Intracellular gallium concentration of *P. aeruginosa* under Ga(NO_3_)_3_ treatment. *P. aeruginosa* were cultured from an initial OD600 of 0.003 at 37°C for 16 hours in the absence or presence of Ga(NO_3_)_3_. The intracellular gallium concentration was determined by an ICP-MS analyzer. Data are presented as mean ± SD. The asterisks represent a significant difference from indicated groups with \*\**P* < 0.01, \*\*\**P* < 0.001 and \*\*\*\**P* < 0.0001 by two-way ANOVA□and□Tukey’s multiple-comparison test.

**Extended data Fig. 6.**
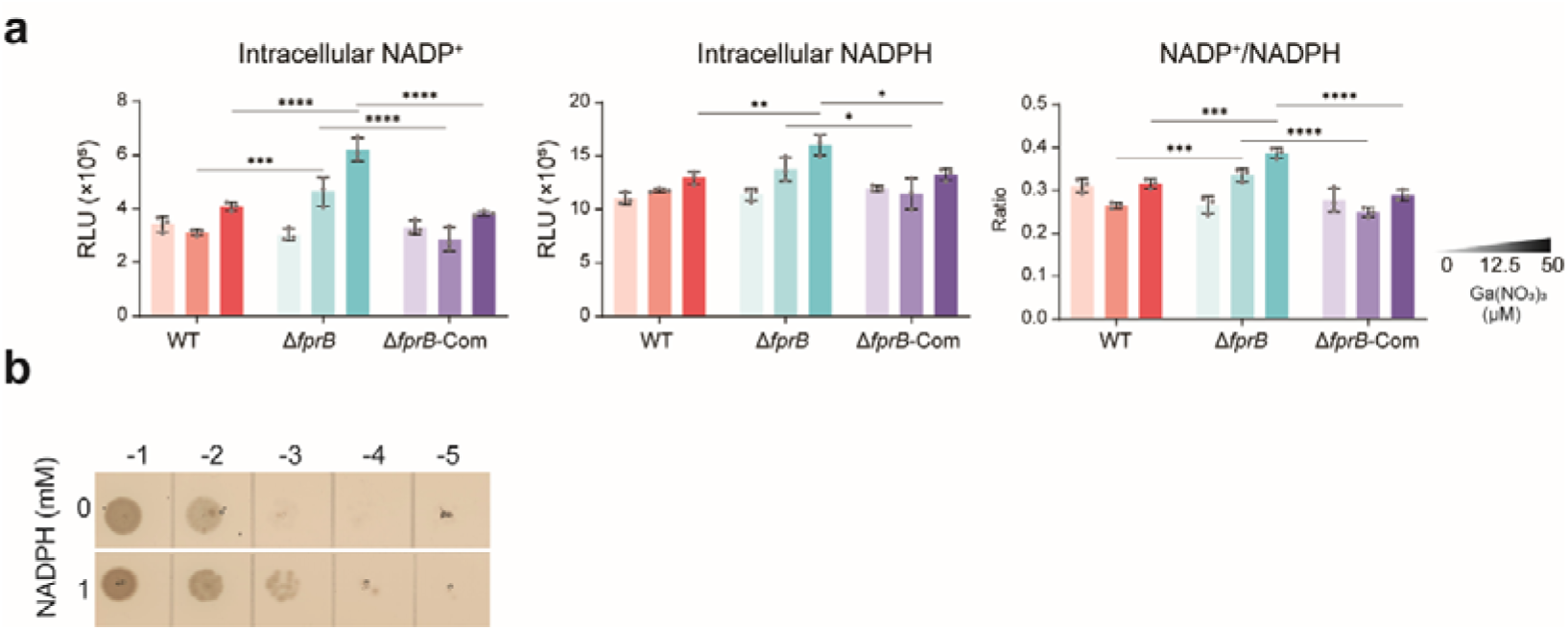
Role of NADP^+^ and NADPH in *P. aeruginosa* growth under Ga(NO_3_)_3_ treatment. **a.** Intracellular NADP^+^ and NADPH accumulation. Bacteria (initial OD600 of 0.01) were cultured for 4 hours in LB broth with or without Ga(NO_3_)_3_. Luminescence value measured by a microplate reader were normalized to final OD600. **b.** Drop plate assay of the Δ*fprB* strain treated with Ga(NO_3_)_3_ and NADPH. Bacterial cultures were adjusted to an initial OD600 of 0.003. Assays were conducted with 6.25 μM Ga(NO_3_)_3_ and/or 1 mM NADPH. Data are presented as mean ± SD. The asterisks represent a significant difference from indicated groups with \*\**P* < 0.01, \*\*\**P* < 0.001 and \*\*\*\**P* < 0.0001 by two-way ANOVA□and□Tukey’s multiple-comparison test.

**Extended data Fig. 7.**
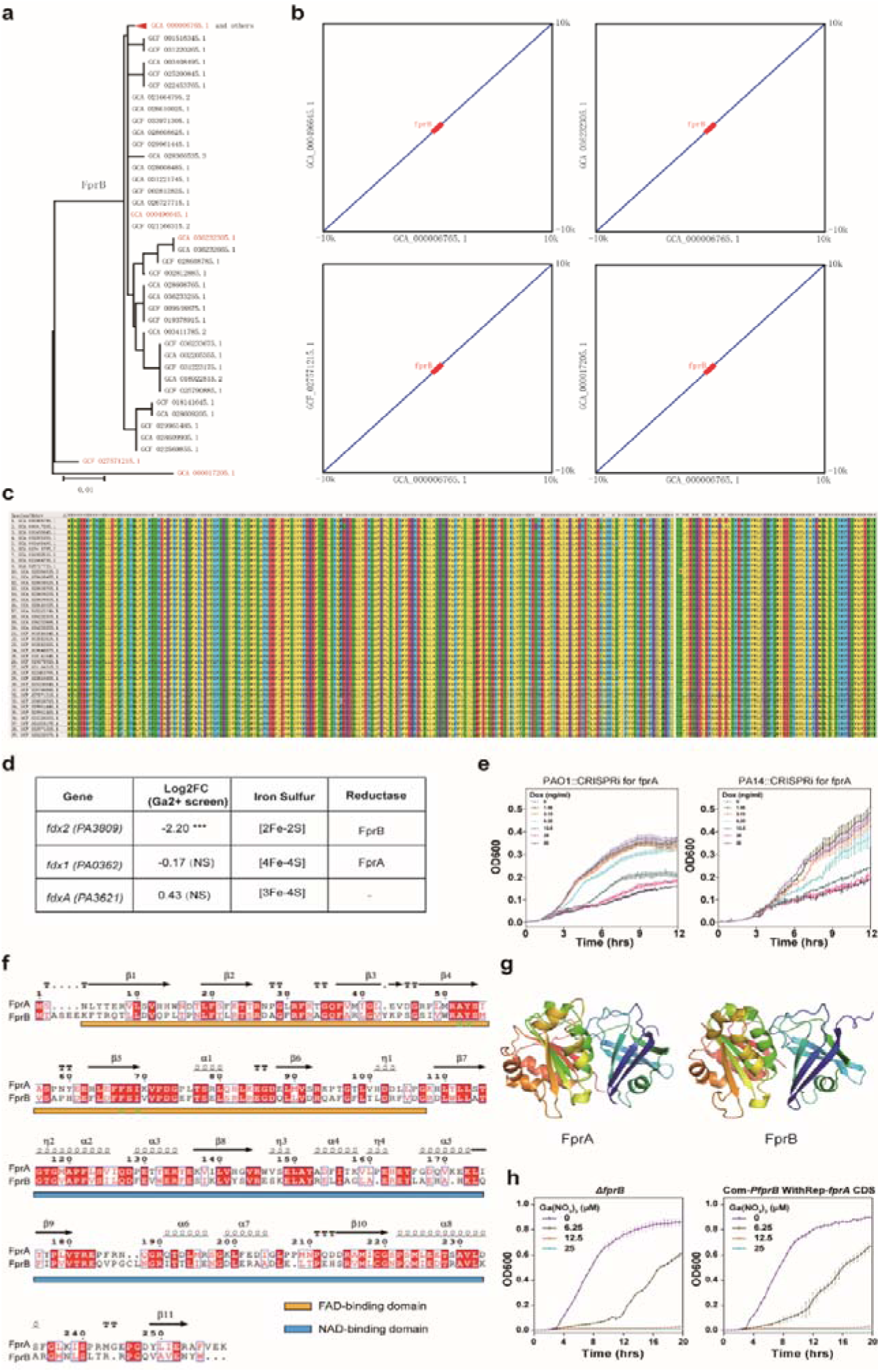
The evolutionary conservation of FprB in *P. aeruginosa*. Analysis of FprB across *P. aeruginosa* strains reveal high conservation at both gene and protein levels. **a.** Neighbor-Joining phylogenetic tree of FprB proteins in representative *P. aeruginosa* strains. In total, 71 phylogenetic sub-clusters were detected from the 981 surveyed *P. aeruginosa* strains. The FprB proteins were collected from the representative strains of each sub-cluster, and used for phylogenetic tree building. The strains for which the genome accessions were highlighted in red were used for further comparative genomic analysis. **b.** Collinearity analysis between *P. aeruginosa* strains representing different FprB sequence clusters for the *fprB* loci (indicated) and their 10k flanking sequences. **c.** Multiple sequence alignment results for the FprB proteins from representative *P. aeruginosa* strains. **d.** CRISPRi-seq screen under Ga(NO_3_)_3_ stress did not identify FprA and its substrate Fdx1 as a contributor to gallium resistance. **e.** Suppression of *fprA* did not significantly increase sensitivity to Ga(NO_3_)_3_. **f, g**. Sequence alignment showing FprA exhibits approximately 42% amino acid sequence identity with FprB in *P. aeruginosa*. **h.** Complementation of *fprA* in the Δ*fprB* mutant did not enhance gallium tolerance.

### Supplementary Tables

Supplementary Table S1. Strains and plasmids used in this work

Supplementary Table S2. Primers used in this work

Supplementary Table S3. gBlocks used in this work

Supplementary Table S4. List of sgRNA and targets in PA14

Supplementary Table S5. PA14 essential genes by CRISPRi-seq

Supplementary Table S6. Classification of genes into five categories by response time and essentiality

Supplementary Table S7. Fitness evaluated by CRISPRi-seq screening under gallium stress

Supplementary Table S8. GO enrichment analysis of CRISPRi-Seq screening for Gallium-treated vs. untreated groups

Supplementary Table S9. Analysis of differentially expressed genes in *fprB* knockout strain vs PAO1 wild type

